# Protein Structure Informed Bacteriophage Genome Annotation with Phold

**DOI:** 10.1101/2025.08.05.668817

**Authors:** George Bouras, Susanna R. Grigson, Milot Mirdita, Michael Heinzinger, Bhavya Papudeshi, Vijini Mallawaarachchi, Renee Green, Rachel Seongeun Kim, Victor Mihalia, Alkis James Psaltis, Peter-John Wormald, Sarah Vreugde, Martin Steinegger, Robert A. Edwards

## Abstract

Bacteriophage (phage) genome annotation is essential for understanding their functional potential and suitability for use as therapeutic agents. Here we introduce Phold, an annotation framework utilising protein structural information that combines the ProstT5 protein language model and structural alignment tool Foldseek. Phold assigns annotations using a database of over 1.36 million predicted phage protein structures with high quality functional labels. Benchmarking reveals that Phold outperforms existing sequence-based homology approaches in functional annotation sensitivity whilst maintaining speed, consistency and scalability. Applying Phold to diverse cultured and metagenomic phage genomes shows it consistently annotates over 50% of genes on an average phage and 40% on an average archaeal virus. Comparisons of phage protein structures to other protein structures across the tree of life reveals that phage proteins commonly have structural homology to proteins shared across the tree of life, particularly those that have nucleic acid metabolism and enzymatic functions. Phold is available as free and open-source software at https://github.com/gbouras13/phold.

## Introduction

Bacteriophages (phages), viruses that infect bacteria, are the most prevalent biological entities on Earth^1^. The persistent growth of metagenomic sequencing data^2^ facilitates the discovery of numerous novel uncultured phages with little or no sequence similarity to known phages^3–7^. Further, increased interest in wet-lab based studies of cultured phages, driven by the resurgence of phage therapy^8,9^ i.e. using phages to target pathogens, particularly those that are anti-microbial resistant, add to our arsenal of relatively well characterised phages^10^.

Despite the increasing number of available phage genomes, phage genome annotation remains extremely challenging. Traditionally, phage genome annotation is conducted using sequence-based homology approaches, consisting of alignment-based methods like MMseqs2^11^ and Diamond^12^, or profile Hidden Markov Model (HMM) based methods like HHblits^13^ and HMMER^14,15^. These algorithms are used by phage genome annotation pipelines^16–20^ to search against general sequence databases like UniProt^21^ or more targeted phage specific databases, like PHROGs^22^ or VOGDB^23^, transferring functional labels to query proteins for strong, likely homologous hits.

However, the divergence and fast evolution of phage genomes limits the ability of sequence-based homology annotation to annotate most phage proteins; over 65% of phage proteins cannot be functionally annotated using sequence-based homology^8^. While protein structure is far more conserved than sequence^24^, until recently it was not possible to utilise structural information for gene annotation due to the small number of solved structures. Driven by the breakthrough of AlphaFold2^25^ in predicting protein structure from sequence, combined with the development of Foldseek^26^ as a rapid protein structural alignment tool fast enough to search through databases containing millions of protein structures to find homology in and below the “twilight zone” of protein sequence alignment (especially below 25% sequence identity)^27^, large-scale structural annotation of phages has recently become possible^28,29^.

This approach has two main bottlenecks. The first is that despite the advent of tools like ColabFold^30^ and ESMFold^31^ that democratise structure prediction, predicting protein structures requires large computational resources even for a single phage. For example, running ColabFold batch with default AlphaFold2 model and parameters for all 66 coding sequences (‘CDS’) of *Escherichia* phage Lambda (GenBank accession: J02459) takes approximately 100 minutes on a NVIDIA A100 GPU, not including multiple sequence alignment (‘MSA’) generation. The second is that while there exist viral sequence databases that cover the general^29,32^ and specific eukaryotic^33,34^ viral spaces, there is no existing database of phage protein structures with high-quality curated functional labels.

Other approaches recently used for genome annotation beyond sequence-based homology are tools that leverage neural networks, particularly protein language models (pLMs). These tools include include EAT^35^ and Gaia^36^ for general annotation and Phynteny^37^, PHANNs^38^, VPF-PLM^39^ and Empathi^40^ for viruses. pLMs are mostly transformer-based language models trained on millions or billions of protein sequences^31,41^ whose resulting embeddings contain a high dimensional understanding of a given protein, including implicitly elements of its structure^42^. However, despite some recent advances^43^, a substantial drawback of using pLM-based methods for gene annotation is the lack of interpretability compared to traditional sequence-based methods, which output sequence alignments and statistical measures of confidence like E-values.

Here we present Phold, a tool for rapid, consistent and accurate annotation of phages using protein structural information. We demonstrate that Phold’s framework, which combines the ProstT5 pLM with Foldseek to search against our new curated database of over 1.36 million mostly phage-derived protein structures, substantially outperforms sequence-based homology methods for phage genome annotation, while maintaining similarly interpretable output.

Applying Phold across the diversity of known cultured phages and uncultured metagenomic viral genomes revealed that over 50% of proteins on an average phage genome can be reliably annotated. We show that Phold using ProstT5 is almost as sensitive as using predicted protein structures but orders of magnitude more resource-efficient, allowing scalability to large metagenome datasets. Finally, by comparing Phold database and other cultivated phage protein structures to protein structures across the tree of life, we show many phage proteins have strong structural similarity to proteins shared across the tree of life, including in *Homo sapiens*.

## Results

### A Comprehensive Database of High Quality and Well-Annotated Predicted Bacteriophage Protein Structures

As a prerequisite for structure-based phage annotation, we first curated and constructed a database of well-annotated phage protein structures. Overall, we predicted structures for 3,166,202 proteins from various sources (hereafter “Phold DB 3.16M”). For the final Phold search database (hereafter “Phold Search DB 1.36M”), we retained a subset of 1,363,704 proteins with high-quality functional annotations (Supplementary Table 1). These include: 441,177 deduplicated Prokaryotic Virus Remote Homologous Groups (PHROG)^22^ proteins from each PHROG cluster containing at least two proteins (i.e. PHROG clusters 1-38880); 70,455 singleton PHROG proteins (i.e. PHROG clusters 38881-109404); 2,205,969 enVhog^44^ proteins (consisting of one representative per enVhog group), of which 562,369 were assigned a PHROG group and kept in Phold Search DB; 392,080 efam^45^ proteins, of which 233,181 were assigned a PHROG group and kept in Phold Search DB; 12,683 diversity-generating retroelement (DGRs) reverse transcriptases^46^ assigned to PHROG 1423; 3,652 anti-CRISPRs (ACRs); 4,804 CARD^47^ antimicrobial resistance proteins; 408 DefenseFinder^48^ proteins; 27,823 Virulence Factor Database (VFDB)^49^ proteins; and 7,152 NetFlax^50^ toxin-antitoxin system proteins. All specialised databases (ACRs, CARD, DefenseFinder, VFDB, NetFlax) were assigned the PHROG category moron, auxiliary metabolic gene and host takeover. All proteins in the Phold Search DB are assigned one of ten general PHROG category labels, along with a more detailed PHROG product label.

Overall, the predicted protein structures were generally of good quality. The mean predicted local distance difference test (pLDDT) for all proteins in Phold DB 3.16M was 75.69 with mean length of 211AA, while the mean pLDDT of the Phold Search DB 1.36M was 82.09 with mean length of 219 AA. The remaining 1.8M structures excluded from the Phold Search DB due to having no assignable PHROG had mean pLDDT of 70.85 and mean length of 207AA. 86.9% of structures included in the Phold Search DB 1.36M had mean pLDDT of at least 70 (Figure 1A). PHROG singleton and enVhog proteins were on average shorter and had lower pLDDT than the other constituents of the Phold Databases (Supplementary Figures 1B, 1C–1H).

**Figure 1.**
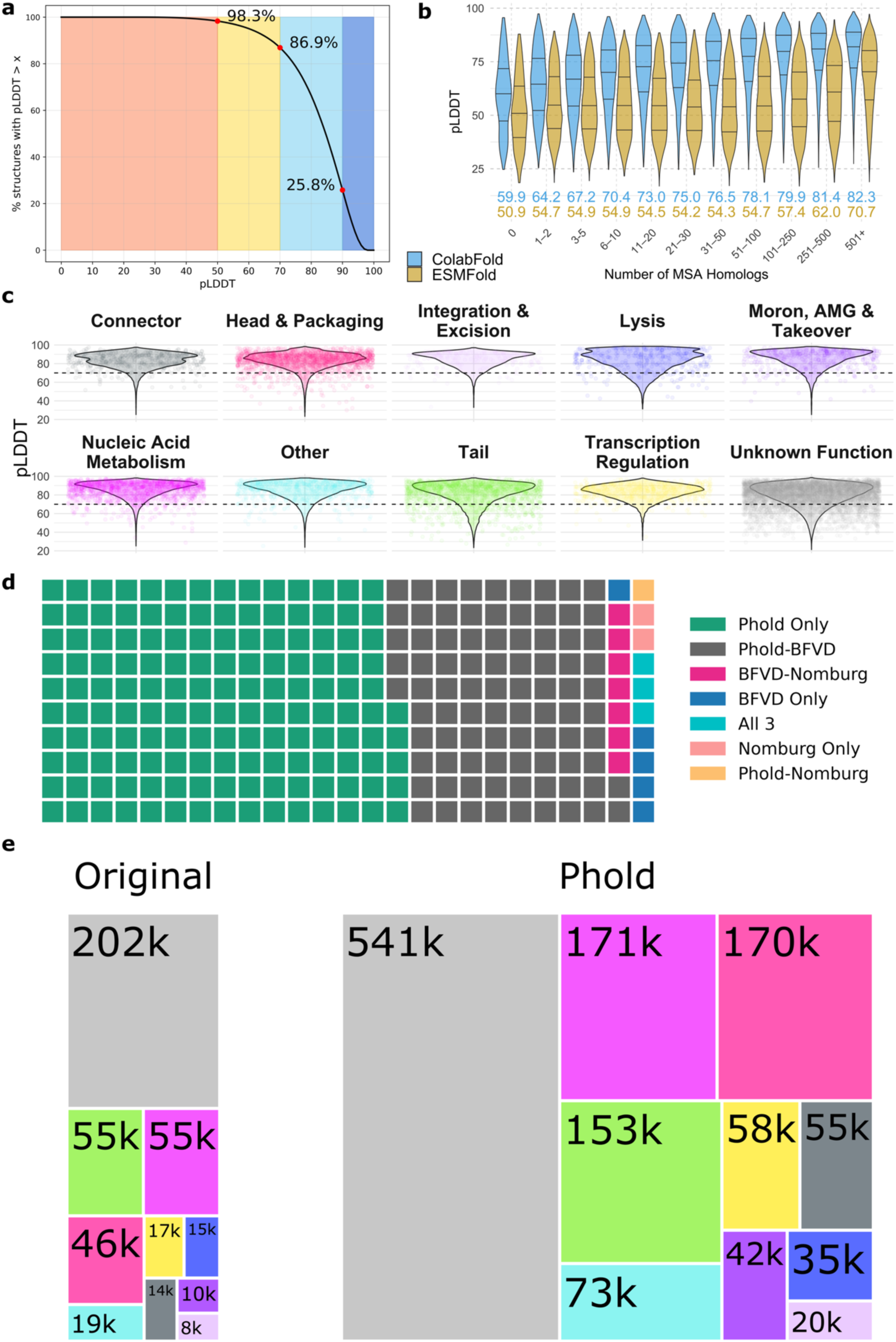
Phold Databases Predicted Quality, Clustering and Annotation Transfer Overview. (a) The proportion of protein structures above a given mean pLDDT value in the Phold Search Database 1.36M (y-axis) plotted against the pLDDT value (x-axis). Colours follow the AlphaFold2 pLDDT colour scheme as implemented in ColabFold’s visualisation. (b) Violin plots showing pLDDT for ColabFold (blue) and ESMFold (gold) (y-axis) against the number of homologs in the protein’s ColabFold MSA (x-axis) in bins for all PHROG, enVhog, efam and PHROG singletons proteins (n=3,099,094), (c) Violin plots showing the distribution of mean pLDDT values for proteins in each of the ten PHROG functional categories. Each protein is a separate point. A subsample of 1% of all proteins were taken for visualisation clarity. (d) Waffle plot showing the number of Foldseek clusters containing exclusively viral proteins (i.e. no AFDB50 members) shared between Phold DB 3.16M, BFVD and Nomburg. One square represents approximately 500 clusters. (e) The number of proteins in each of the ten PHROG functional categories in the original PHROGs v4 database (left) compared to the Phold Search DB (right). The colour scheme matches (b), with the largest grey boxes constituting unknown function proteins belonging to a PHROG group.

We predicted structures for all structures using both ColabFold and ESMFold, selecting the structure with the highest pLDDT score (Supplementary Note 1). ColabFold outperformed ESMFold for the majority (2,752,972/3,166,202: 86.5%) of proteins, with an overall higher mean pLDDT (74.8 vs 61.2). We observed that while structure quality was correlated with MSA depth for both methods, for ESMFold proteins with fewer than 100 homologs in the ColabFold MSA, there was no correlation between ESMFold pLDDT and number of homologs with generally low-quality predictions (Spearman’s ρ: -0.01, median pLDDT: 54.6). For those with at least 100 homologs, there was a positive correlation and more confidently predicted structures (Spearman’s ρ: 0.30, median pLDDT: 67.6) (Figure 1B; Supplementary Figures 2A–C).

For PHROG singletons and enVhogs, we enriched the MSAs by extending ColabFold’s default search databases to include our collection of 129,944,764 viral proteins. For PHROG singletons, we produced and compared ColabFold predictions with and without enriched MSAs. Enrichment increased the number of homologs from a median of 19 to 53, improving pLDDT for 41,802/70,524 (59.3%) singletons. The median ColabFold pLDDT increased from 74.4 (base MSA) to 76.4 (enriched), yielding an overall median ColabFold pLDDT of 77.3 when we took the highest pLDDT from the two methods. The number of high-quality (pLDDT >70) predictions increased by 4,406/70,524 (6.2%) (Supplementary Figure 3A). Proteins with base MSAs that were neither completely empty nor very deep and high quality had the largest improvement in pLDDT with the additional sequences (base MSA homologs from 11-20 improved by median 1.22 pLDDT; Supplementary Figures 3B–C).

Overall, 29,171/38,880 (75.0%) of PHROG groups containing 1,180,385/1,319,865 (89.4%) proteins had mean pLDDT of at least 70. The mean pLDDT for all nine functional PHROG categories was high, ranging from 81.1 for tail proteins up to 85.6 for nucleic acid metabolism proteins (Supplementary Table 2). Poor predictions were more commonly generated for tail and lysis category proteins compared to the other categories (Figure 1C), with 10^th^ percentile pLDDT means of 63.5 and 66.4, respectively, compared to a minimum of 71.4 for any other functional category (Supplementary Table 2). The quality of predicted structures for PHROGs with unknown function was high (mean pLDDT 79.8) and much higher than enVhog and efam proteins with no assignable PHROG (mean pLDDT 70.2) (Supplementary Figure 1). 815/856 unique PHROG product descriptions had mean pLDDT of at least 70 (Supplementary Table 3).

### Phold’s Database Expands the Known Phage Protein Structure Space

To understand the novelty of Phold’s predicted viral protein structures, we co-clustered Phold DB 3.16M with AlphaFold Database’s AFDB50 representatives^51,52^ (n=53,665,860) and two existing viral protein structure databases–BFVD^29^ (n=351,242) and Nomburg^33^ (n=74,129)– with Foldseek using a 70% coverage cutoff. 57,257,433 proteins yielded 16,590,479 clusters, 14,395,143 of which were singletons (25.1%), while 42,862,290 (74.9%) belonged to 2,195,336 clusters with at least two members. Remarkably, 441,724 clusters summarizing 23,633,190 proteins contained at least one member from any viral database (i.e. Phold DB 3.16M, BFVD or Nomburg; ‘viral clusters’) (Supplementary Table 4). 250,319 (56.7%) of these viral clusters contained both AFDB50 and Phold DB proteins only (Supplementary Figure 4; Supplementary Tables 4–5). Phold DB proteins were present in 420,656 viral clusters containing a total of 22,972,540 members (95.2% of non-singleton viral clusters containing 97.2% of non-singleton viral cluster members), while AFDB50 proteins were present in 314,971 viral clusters containing 23,210,560 members (71.3% of non-singleton viral clusters containing 98.2% members). 73,525 clusters containing 215,138 members (16.7% of clusters containing 0.9% of members) were unique to Phold DB (Supplementary Figure 4; Supplementary Tables 4–5), suggesting Phold DB primarily expands the diversity of less common viral structures and folds.

Of the 441,742 viral clusters, 126,753 clusters containing 422,630 proteins did not contain any AFDB50 member. Of these 126,753 clusters, Phold DB structures were present in 119,875 (94.6%), with 73,525 containing only Phold DB proteins (58.0%) (Figure 1D). There was substantial overlap between Phold DB and BFVD, with 44,278 (35.0%) clusters containing only Phold DB and BFVD proteins. This suggests that despite its relatively small size (i.e. almost ten-fold smaller than Phold DB), BFVD covers a substantial portion of viral protein structure space outside of the AFDB.

Overall, 54.5% (1,957,655/3,591,573) of viral database proteins could be structurally clustered with AFDB50. This suggests that despite explicitly omitting viral proteins, the AFDB implicitly includes proteins with homology to many viral and phage protein structures, covering a substantial portion of the known viral protein structure space. This could be due to the inclusion of provirus protein structures present in AFDB, particularly those derived from prophages^53^. Additionally, it may explain why our efforts to fine-tune ProstT5 using Phold DB protein structures showed only modest improvements compared to using the standard ProstT5 (Supplementary Note 2; Supplementary Figure 5; Supplementary Table 6).

### Structural Similarity Expands Annotation Labels in Phold’s Database

The major impact of the original PHROG database was the manual high-quality expert-curated annotations provided for 5,133/38,880 PHROGs (13.2%) containing 239,059/440,550 unique proteins (54.3%), along with 165/70,234 PHROG singletons (0.2%). Given the structural similarities between Phold DB proteins with the AFDB and BFVD, we used structural similarity searches to guide manual curation (see Methods) to transfer functional annotations to an additional 2,798 PHROGs containing 30,490/440,550 (6.9%) PHROG proteins along with 3,019 singletons (Supplementary Figure 6; Supplementary Table 7). Combining proteins enVhog and efam proteins assigned PHROGs and specialised databases; 822,262 unique proteins in the Phold Search DB are assigned functional annotations, while a further 541,442 are included with a PHROG but with yet unknown function (Figure 1E).

### Phold’s Framework for Phage Protein Annotation

Phold combines the structure-informed pLM ProstT5^54^ with the structure-comparison tool Foldseek^26^ to search against the Phold Search DB 1.36M containing 1,363,704 protein structure predictions. Phold’s workflow consists of four stages: (a) initial gene calling and genome feature annotation; (b) Foldseek 3Di token prediction using ProstT5 or 3Di token extraction from pre-computed PDB/mmcif; (c) Foldseek structure comparison and annotation transfer; and (d) plotting and summary generation (Figure 2).

**Figure 2.**
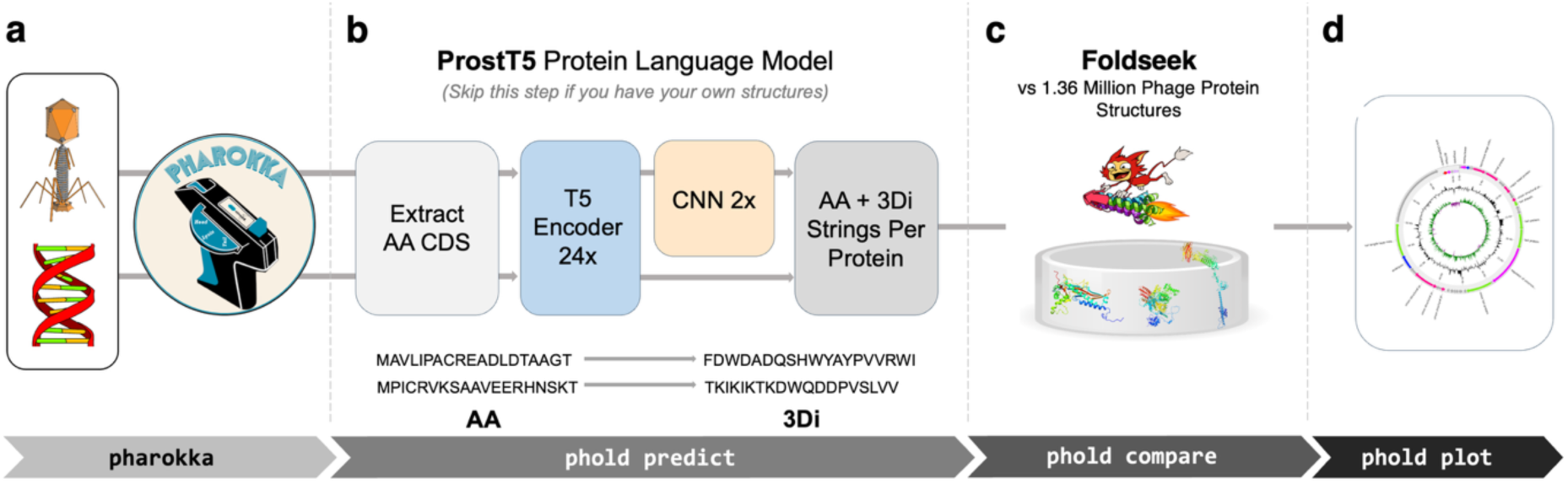
Phold’s workflow for annotating bacteriophage genomes. (a) Phold begins with assembled phage genomes. Pharokka is run first to predict coding sequences (CDS) and other genomic features such as tRNA and tmRNAs and to annotate CDS using sequence-based homology-detection methods. (b) The ‘phold predict’ module predicts Foldseek 3Di tokens for each residue of each CDS using the ProstT5 pLM encoder and CNN. Alternatively, predicted protein structures can be used instead of this module if available. (c) The ‘phold comparè module then compares every CDS using the amino acid and ProstT5 predicted 3Di tokens against the Phold Search DB 1.36M containing 1,363,704 protein structures. The top-ranking database hit (based on Foldseek E-value) with a non-hypothetical function is assigned to each CDS. (d) Summaries and genomic plots can then be created with the ‘phold plot’ module.

Phold accepts Pharokka^16^, Bakta^55^ or NCBI GenBank formatted genomes as the desired input. Alternatively, Phold can take nucleotide FASTA as input and utilises Pyrodigal-gv^18,56,57^ as a gene caller, as it has been shown to improve the annotation of phages that use alternative genetic codes^58^. Phold extracts all predicted coding sequences (CDS), generates 1024-dimensional embeddings for each CDS residue using the 1 billion parameter ProstT5 pLM encoder. These embeddings are then fed into a 2-layer convolutional neural network (CNN) trained to predict the Foldseek 3Di token at each residue. Alternatively, 3Di tokens are extracted from predicted protein structures in the PDB or mmCIF format if available. Phold then uses Foldseek to search against Phold’s database, which consists of predicted phage protein structures based on the PHROGs database, along with other specialised sub-databases. The top-ranking hit below the E-value with an assigned function in the database is finally transferred to annotate each query protein.

### ProstT5 Confidence is Well Calibrated and Positively Correlated with Protein Structure Quality

We next analysed reliability of generating ProstT5 3Di tokens compared to established methods (i.e. from ColabFold/Alphafold2 structures) and reliability metrics (i.e. pLDDT) using the ProstT5 confidence metric (Supplementary Note 3; Supplementary Figures 7–8). Across all proteins in Phold DB 3.16M, we found that the empirical similarity between ProstT5 3Di predictions and 3Di tokens generated from predicted protein structures was strongly positively correlated (Spearman’s ρ = 0.87) (Figure 3A). ProstT5 confidence was also positively correlated with mean pLDDT (Spearman’s ρ 0.40) (Figure 3B). ProstT5 confidence, though having the same range of possible values as pLDDT (i.e. 0-100), was generally substantially lower overall (mean 58.96 compared to 75.69 for pLDDT). ProstT5 confidence was higher for the Phold Search DB 1.36M proteins (mean 61.33) compared to the proteins without PHROG group assignment (mean 57.17). Although correlated, we caution interpreting ProstT5 confidence exactly like pLDDT, particularly in terms of any hard cutoffs (i.e. the widely used pLDDT 70 threshold does not translate to a specific ProstT5 confidence score).

**Figure 3.**
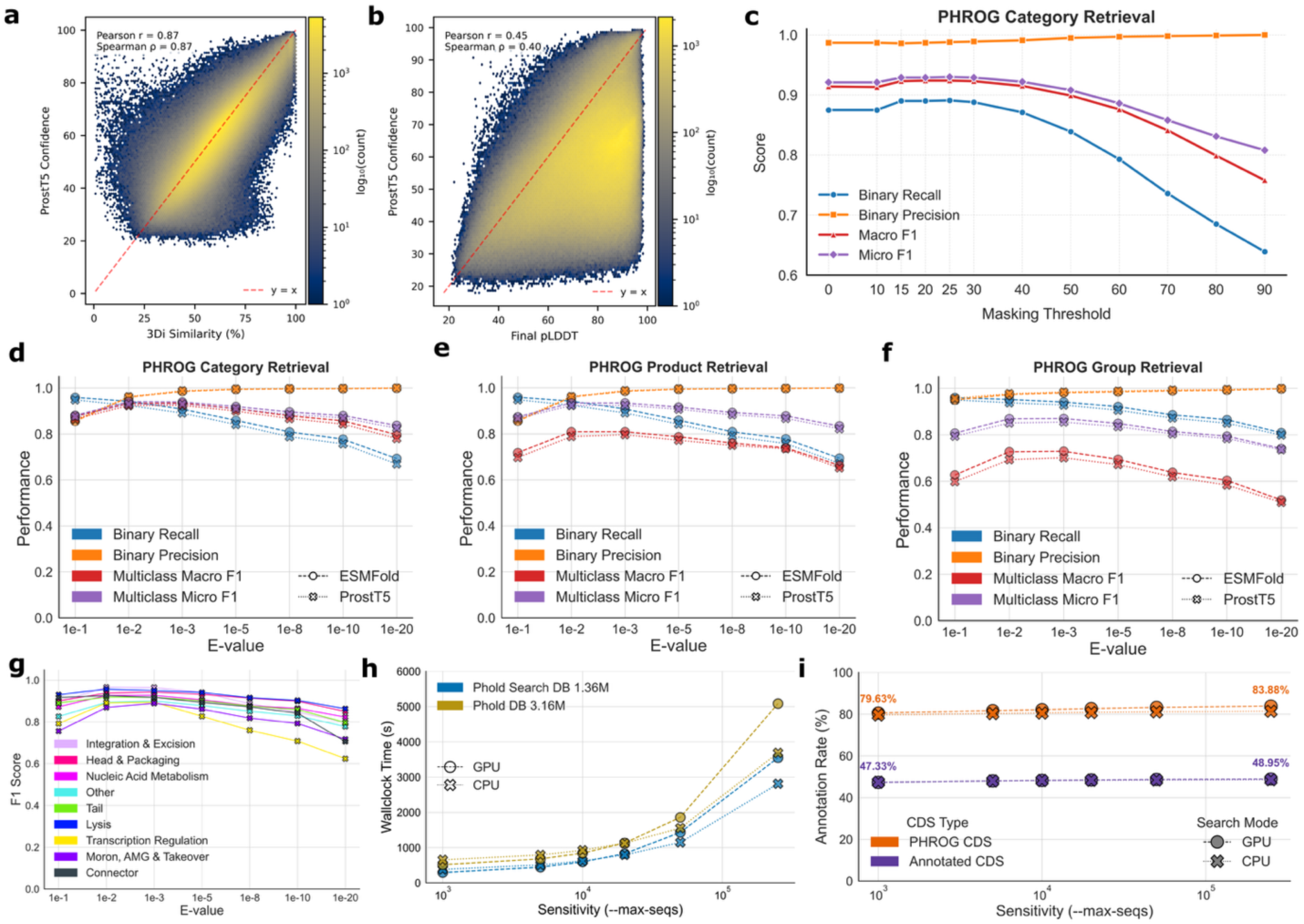
Phold with ProstT5 annotation performance (a) Empirical sequence similarity between ProstT5-CNN generated 3Di tokens and the Foldseek VQ-VAE derived 3Di tokens (x-axis) and ProstT5 confidence (y-axis) for all Phold DB 3.16M protein structures. (b) Final pLDDT (i.e. the higher of ColabFold or ESMFold pLDDT) (x-axis) and ProstT5 confidence (y-axis) for all Phold DB 3.16M protein structures. (c) PHROG category annotation performance of Phold at E-value=0.001 when masking low ProstT5 confidence residues below threshold (x-axis) on the INPHARED 182 dataset compared to the pseudo ground truth of annotations with ColabFold generated structures at E-value 0.001. Binary recall and precision measure whether ProstT5 with masking can annotation any non-hypothetical annotation for annotated ground truth proteins, while multiclass F1 scores are measured across all 9 PHROG categories. (d) PHROG category annotation performance of Phold using ProstT5 (crosses) and ESMFold (circles) at a variety of E-values on the INPHARED 182 compared to the pseudo ground truth of annotations with ColabFold generated structures at E-value 0.001. (e,f) The same but for the PHROG product annotation task (i.e. more specific annotations) (e) and PHROG group assignment task (f). (g) Individual PHROG category annotation performance of Phold using ProstT5 at a variety of E-values on the INPHARED 182 compared to the pseudo ground truth of annotations with ColabFold generated structures at E-value 0.001. (h) Runtime of ‘phold comparè on the INPHARED 182 dataset (y-axis) for different sensitivity values (i.e. Foldseek ‘--max-seqs’ parameter that dictates the number of hits that pass through the prefilter) (x-axis), using both Foldseek-CPU and Foldseek-GPU and Phold Search DB 1.36M (blue) and Phold DB 3.16M (gold). (i) Functional (purple) and PHROG (orange, i.e. also including unknown function PHROG hits) annotation rates with Phold using ProstT5 across the 16,460 CDS in INPHARED 182 (y-axis) using the Phold Search DB 1.36M for different sensitivity (i.e. Foldseek ‘--max-seqs’) values (x-axis).

### Phold With ProstT5 Is Orders of Magnitude More Resource Efficient Than with ESMFold or AlphaFold2 While Yielding Similar Annotation Performance

We next benchmarked the annotation performance of Phold using ProstT5-based 3Di predictions with Foldseek. We chose four benchmarking datasets, ranging from relatively well-studied isolated and taxonomically classified phages (INPHARED^10^ 182) to metagenomically assembled phage genomes from various sources (Tara^59^, Cook^3^ and Crass^6^). The benchmarked task was whether Phold with ProstT5 could annotate the PHROG category (general, practically useful), product (specific, practically useful), or PHROG group (very specific, less practically useful) for each protein compared to using the pseudo ground truth of annotation using Foldseek with ColabFold (AlphaFold2) structures top hits at an E-value cut-off of 0.001.

We first benchmarked the impact of masking residues with low ProstT5 3Di prediction confidence at different thresholds to see whether masking may be used as an estimate of 3Di prediction quality akin to pLDDT for protein structure prediction. Masking residues below 15, 20, 25 and 30 ProstT5 confidence led to improvement in recall with minimal impact on precision on INPHARED 182, with the binary annotation F1 increasing from 0.927 to 0.937 and multiclass micro-averaged F1 from 0.921 to 0.930 compared to no masking (Figure 3C). For the specific PHROG product task, masking residues below 25 increased multiclass micro-averaged F1 from 0.916 to 0.924 (Supplementary Figure 9A), with similar results for the general PHROG category prediction task (multiclass micro F1: 0.862 vs 0.878) (Supplementary Figure 9B). When masking residues above ProstT5 confidence of 30, performance decreased, driven by collapsing recall (Figure 3C; Supplementary Figure 9; Supplementary Tables 8–9) as increasingly more 3Di residues were masked. Similar results were observed for the three metagenomic phage datasets (Supplementary Figure 10). We chose to implement a default ProstT5 confidence masking threshold of 25 in Phold.

We then benchmarked Phold using ProstT5 versus ESMFold-derived structures across varying Foldseek E-value thresholds, using pseudo ground truth from ColabFold (E=0.001). ProstT5 enabled ∼50–100× faster 3Di token generation than ESMFold (Supplementary Table 10), with only marginal reductions in annotation performance across all four datasets (Figures 3D–F; Supplementary Figures 11–12; Supplementary Tables 11–12). On the INPHARED 182 dataset of relatively well-characterised phages, both methods performed similarly across PHROG group, product, and category levels, with optimal precision-recall trade-offs at E=0.01–0.001. Phold with ProstT5 showed strongest performance on the more practically useful broader annotation tasks (F1=0.939 for category; F1=0.924 for product), with reduced accuracy at the group level (F1=0.855 at E=0.001). Performance was robust across all nine PHROG categories on INPHARED 182 (F1=0.889–0.965; Fig. 3G), and generally consistent for Cook and Tara datasets, but lower on Crass, particularly for the uncommon connector category (F1=0; *n*=11; Supplementary Figure 12B), indicating the difficulty of this dataset. Notably, the therapeutically relevant integration and excision category showed high performance across datasets (F1=0.872–0.965).

As Foldseek search sensitivity can be varied and run with either CPU- or GPU-acceleration^60^ applying different prefilter algorithms, we next ran a series of ablations to test the impact of varying hardware and sensitivity and database pre-clustering on Phold’s performance and runtime (Supplementary Table 13). Using INPHARED 182, Phold using ProstT5 with Foldseek-GPU was faster than Foldseek-CPU at default sensitivity. Annotation performance increased marginally with sensitivity (Figure 3H), at the cost of substantial runtime increases (Figures 3I). Phold with Foldseek-GPU was able to detect more CDS hits to any PHROG (i.e. including PHROGs with unknown function) compared to Foldseek-CPU regardless of sensitivity (Figure 3I), but there were minimal differences in terms of functionally annotated proteins (Figure 3I). This trend is observed because Foldseek-GPU performs full alignment of all sequences while its CPU counterpart performs *k*-mer-based pre-filtering first and only aligns high-quality hits. Pre-clustering the Phold Search DB did not substantially impact annotation rate or runtime (Supplementary Note 4; Supplementary Figure 13).

### Phold Enables Consistent Annotation of Over 50% of Genes for an Average Phage Genome

We next compared Phold’s annotation performance with that of existing sequence homology-based phage annotation methods. We compared Phold to Pharokka for several reasons: Pharokka is widely used in the phage genome annotation field; it has options to search using either position-specific scoring matrices (PSSM) profiles (MMseqs2) or with HMM (PyHMMER) based homology inference; and it utilises the same hierarchy of PHROG categories and product annotation labels, making it easy to compare functional labels across different homology detection methods.

We tested the annotation performance on 1,419 INPHARED viruses from distinct genera (see Supplementary Table 14 for information on all viruses included). We compared four methods: Pharokka using MMSeqs2, Pharokka using PyHMMER, Phold with Foldseek and ProstT5 for 3Di and Phold with Foldseek and 3Di from ColabFold predicted structures. The 1,419 viruses contained 148,194 CDS. We generated ColabFold structure predictions for 147,946 proteins, consisting of all but the longest proteins (over 3,000AA) (mean pLDDT of 77.05). This dataset represents the largest consistently annotated set of protein structural predictions across the diversity of known bacteriophage and archaeal virus genomes.

Phold with ProstT5 annotated 73,268 (49.4%) CDS, increasing to 76,322 (51.5%) using ColabFold-generated structures, a marked increase from Pharokka’s 51,399 (34.7%) with MMseqs2 and 55,923 (37.7%) with PyHMMER. Phold was able to improve annotations for all 9 PHROG categories (Figure 4A), with transcriptional regulation showing the highest proportional increase (1,742 with Pharokka with MMseqs2 vs 3,323 Phold with structures) and lysis the lowest (2,879 vs 3,847). The annotation improvements across PHROG categories for the Cook, Tara and Crass metagenomic datasets were similar (Supplementary Figure 14A–C).

**Figure 4.**
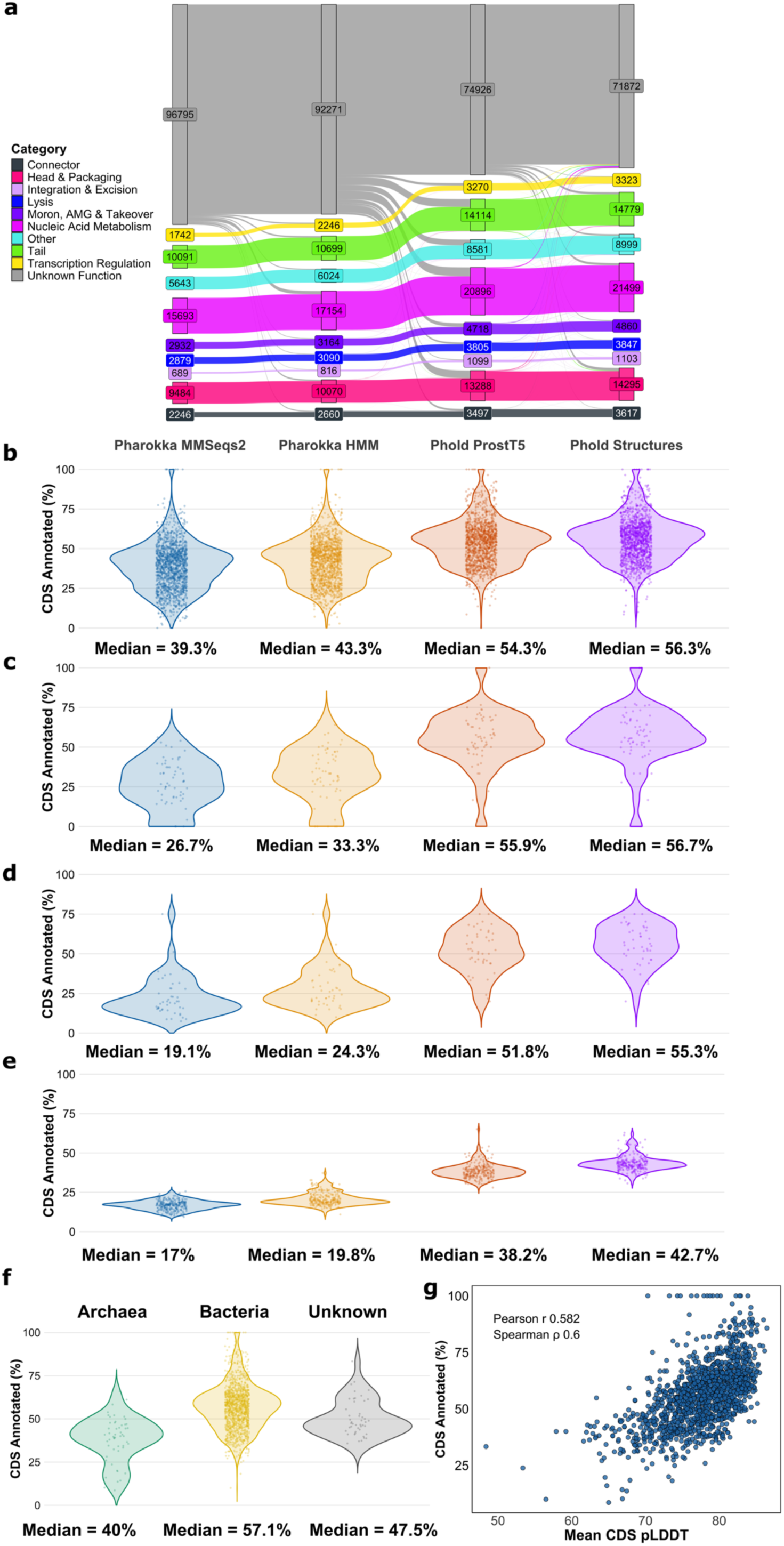
Phold annotates over 50% of coding sequences (CDS) in the average phage genome. (a) Flow of annotations for all ten PHROG categories (including unknown function) between the four annotation methods: Pharokka with MMSeqs2 (left); Pharokka with HMMs (PyHMMER) (centre-left), Phold with ProstT5 (centre right) and Phold with ColabFold structures (right). (b-e) Percentage of CDS annotated (y-axis) for every virus in the (b) INPHARED 1419 (c) Cook (d) Tara and (e) Crass datasets for each of four annotation methods. Each point represents a phage, with the median indicated at the bottom of each plot. (e) Percentage of CDS annotated (y-axis) for every virus in the INPHARED 1419, grouped by the domain of the host taxa. Unknown indicates that the virus was metagenomically assembled, or the host was not available in the metadata associated with the genome. (f) Mean CDS ColabFold pLDDT (x-axis) against the percentage of CDS annotated (y-axis) for every virus in INPHARED 1419. Each point represents the mean CDS pLDDT for one virus.

The median per-phage annotation rate for Phold was 54.3% with ProstT5 and 56.3% with structures, similar to previously reported annotation rates for phages using Foldseek with ColabFold structures searching against BFVD^29^ or AFDB^51^ (Figure 4B). Phold annotation rates were similar for Cook (Figure 4C) and Tara (Figure 4D), despite having lower Pharokka annotation rates. While Phold annotation rates were lower for Crass, they were still more than double Pharokka’s (Figure 4E). These results indicate that Phold is especially useful compared to sequence-based methods for annotating more distant, difficult-to-annotate phages with computation efficiency allowing scaling to large metagenomic-scale datasets. On a single NVIDIA A100 40GB GPU, ProstT5 3Di inference for all 1419 INPHARED viruses with Phold took 3942 seconds, whereas generating large-scale protein structure predictions with ColabFold/AlphaFold2 involves generating MSAs, requires large-scale computational infrastructure, and is three-to-four orders of magnitude more resource intensive^54^.

1393/1419 (98.2%) and 943/1419 (66.5%) of the INPHARED viruses had at least 30% and 50% of their CDS annotated respectively. Archaeal viruses had a lower annotation rate (median 40% *n*=52) compared to bacterial viruses (median 57.1% *n*=1319) and with unknown host (median 47.5% *n*=48) (Figure 4F). Of the host genera with at least ten infecting viruses, the median annotation rate ranged from 38.3% (*Halocarcula*) to 69.3% (*Burkholderia*) (Supplementary Figure 15A). Phages for *Staphylococcus* and *Escherichia* had among the highest annotation rates (67.9% and 63.2%). The percentage of Phold annotated CDS was strongly correlated with the mean ColabFold pLDDT for all CDS in the virus, regardless of whether ProstT5 or ColabFold structures were used (Phold with ColabFold structures: Spearman ρ 0.60 Figure 4G: Phold with ProstT5: Spearman ρ 0.584 Supplementary 15B). This is unsurprising given the dependence of both annotation and structure prediction quality on the existence of homologous proteins.

### Phold Allows for Large Scale Hypothesis Generation of Unknown Phage Protein Functions

Detailed analysis of the 1419 INPHARED genome annotations revealed an interesting wealth of annotations to the specialised databases employed by Phold. 5/1419 phages contained six proteins with homology to CARD antimicrobial resistance proteins, five of which have sequence similarity below 30%. For instance, *Rhizobium* phage RHph_Y65 (GenBank accession: NC_070967) possesses a CDS annotated by Phold with high structural similarity (TM-score 0.93, LDDT 0.76) to the CARD trimethoprim resistant dihydrofolate reductase gene DfrA37 with sequence similarity of only 26% (Figure 5A), while *Vibrio* phage VAP7 (GenBank accession: NC_048765) also possesses a CDS with high structural similarity (TM-score 0.91, LDDT 0.77) to the related CARD trimethoprim resistant dihydrofolate reductase gene dfrA26 (Supplementary Figure 16A). Both hits were also found with Phold ProstT5 (Foldseek E-values 1.8E^-17^, 5.3E^-18^), but missed with Pharokka using MMseqs2-based sequence homology searches against CARD. The presence of phage-encoded dihydrofolate reductases does not always confer trimethoprim resistance^61^, but it has been shown to occur in some prophages^62^, and has been implicated in resistance phenotypes.

**Figure 5.**
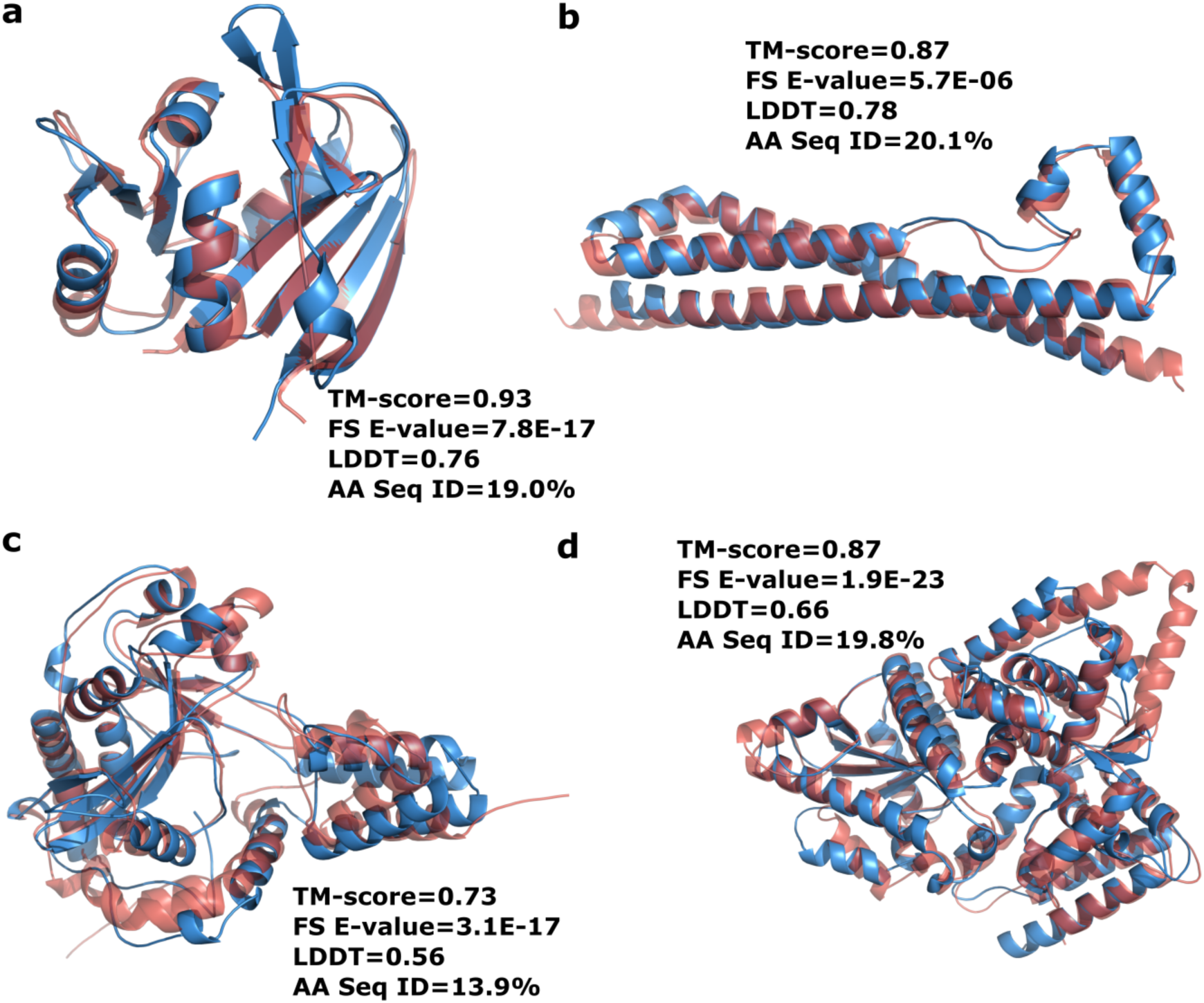
Phold allows for the discovery of phage proteins with high structural similarity to anti-microbial resistance and anti-phage defense proteins. (a) ColabFold predicted structures of a predicted dihydrofolate reductase from *Rhizobium* phage RHph_Y65 (GenBank accession: NC_070967) (blue) and CARD trimethoprim resistant dihydrofolate reductase gene DfrA37 (red) (b) Predicted cyclase from *Clostridioides* phage phiCD27 (GenBank accession: NC_011398) (blue) and the Pycsar defense system cyclase (red) (c) Predicted DruA-like protein from *Synechococcus* phage S-PRM1 (GenBank accession: NC_055761) (blue) and DruA protein from the Druantia Type I defense system (red) (d) Predicted GAPS1-like protein from *Klebsiella* prophage ST437-OXA245phi4.1 (blue) and GAPS1 defense system protein (red). “FS E-value” = Foldseek E-value. “AA Seq ID” = amino acid sequence identity.

Proteins with hits to Anti-CRISPRs were found in 82/1419 INPHARED viruses containing 112 predicted Anti-CRISPRs, while 117 proteins with structural similarity to virulence factors were found in 85/1419 viruses. 62 proteins with structural similarity to the DefenseFinder database of bacterial anti-phage proteins were found in 52/1419 viruses. For example, two *Clostridioides* prophages (phiC2 GenBank accession: NC_009231 and phiCD27 GenBank accession: NC_011398) encode a CDS with only 20% sequence identify to the Pycsar defense system^63^ cyclase despite extremely high structural similarity (TM-score 0.87, LDDT 0.77) (Figure 5B), while six *Synechococcus* phages encode CDS with very low sequence identity (12.7-17.1%) but high structural similarity to the DruA protein from the Druantia Type I defense system^64^ (TM-score 0.73, LDDT 0.56) (Figure 5C). Additional examples include a CDS from *Klebsiella* prophage ST437-OXA245phi4.1 (GenBank accession: NC_049448) with similarity to the GAPS1 system^65^ (TM-score 0.87, LDDT 0.66) (Figure 5D), AriaA homologs from both subsystems of the PARIS defense system^66^ on *Fusobacterium* phage vB_FnuS_FNU3 (Genbank accession: OQ808965) and *Yersinia* phage vB_YenM_31.17 respectively (Supplementary Figure 16B–C) and a letS protease homolog to the Gao_let defense system^67^ on *Enterocbacter* phage phiT5282H (GenBank accession: NC_049429) (Supplementary Figure 16D).

The prevalence of homologs to bacterial anti-phage defense system proteins in the 1419 INPHARED viruses emphasises that phages may serve as a reservoir of anti-phage defence systems that are prophage encoded^68^, a concern that supports the exclusion of using prophages in therapeutic contexts^69^. It also suggests that phages may use similar mechanisms to defeat or mimic known bacterial defense systems in the phage-host arms-race^70^.

### Many Phage Proteins Have Structural Similarity to Proteins Across the Tree of Life

The scale and quality of protein structure predictions and annotations of the Phold DB 3.16M and the 1419 INPHARED viruses enables broad scale comparison to proteins belonging to vastly different taxa^71,72^. Using thresholds of 70% bidirectional coverage, a maximum Foldseek E-value of 0.01 and minimum TM-score of 0.6 to ensure a high likelihood of a shared fold across the protein^73^, we found 346,495 (10.9%) of Phold DB 3.16M proteins had AFDB hits to *Eukaryota* proteins, 632,793 (20.0%) to *Archaea* proteins and 1,176,208 (37.1%) to *Bacteria* proteins. 296,912 (9.4%) of proteins had hits to all three domains of life (Supplementary Table 15). For INPHARED 1419, the numbers were proportionally similar between the three domains and higher overall (24,654 (16.7%) for *Eukaryota*, 43,155 (29.2%) for *Archaea* and 74,103 (50.1%) for *Bacteria* and 22,129 (15.0%) with hits to all three domains) (Supplementary Table 16). Proteins with metabolic or enzymatic functions belonging to nucleic acid metabolism, other, and moron, auxiliary metabolic gene and host takeover PHROG categories had the highest proportion of hits (Figures 6A–B). Transcription regulation and unknown function categories had the lowest proportions of hits (Figures 6A–B), indicating that finding structural similarity to proteins in these PHROG categories is potentially more difficult due to their shorter length^74^ (Supplementary Table 1) and that proteins in these categories (especially unknown function) are more likely to be unique to phages.

**Figure 6.**
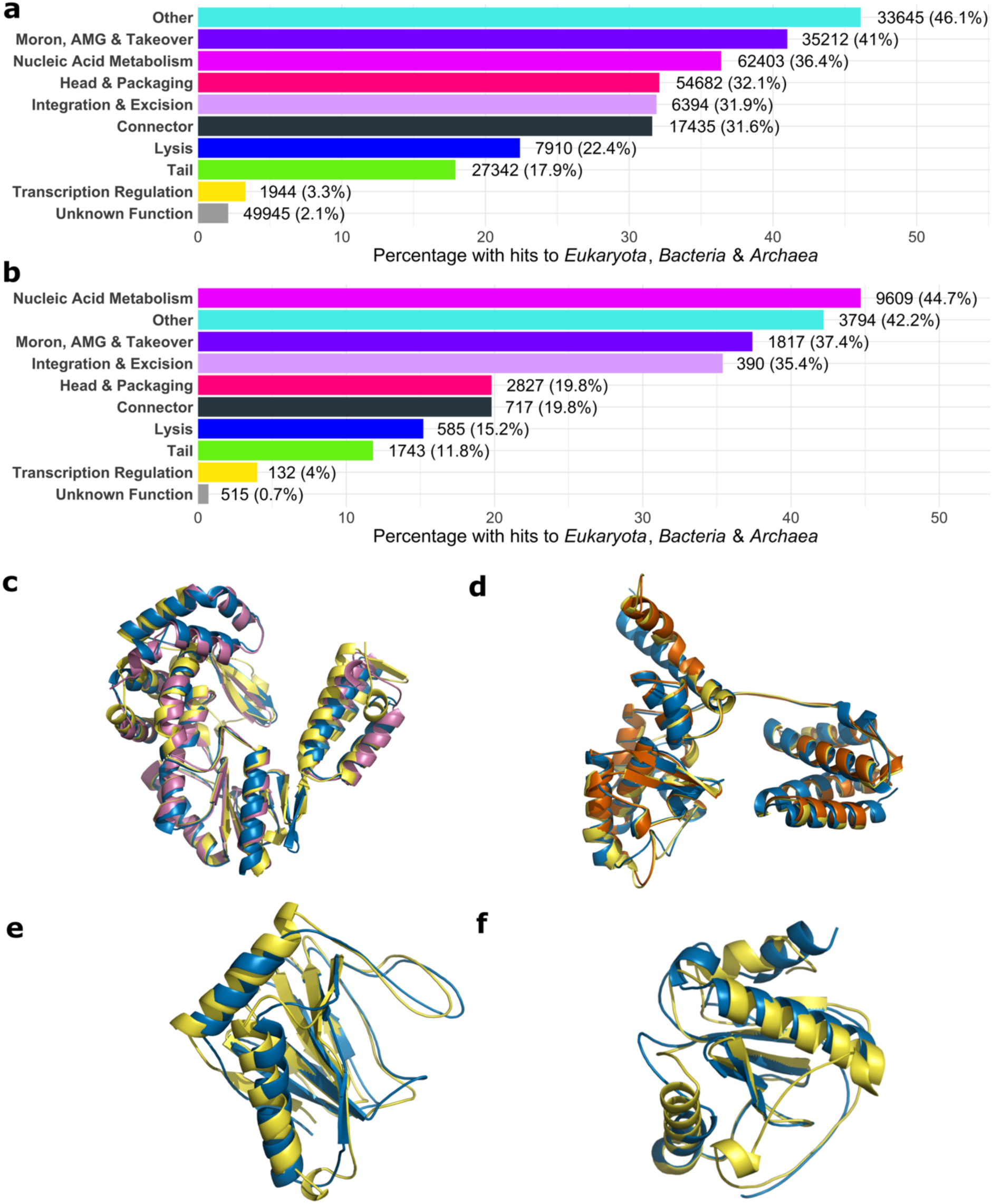
Structural similarity of phage proteins across the tree of life. Percentage of proteins belonging to each PHROG category that have strong structural similarity (Foldseek E-value < 0.01 and alignment TM-score > 0.6) to all three domains of life (*Eukaryota*, *Bacteria,* and *Archaea)* from AlphaFold Database proteins for (a) Phold DB 3.16M proteins and (b) INPHARED 1419 proteins. The raw number and percentage per category are indicated to the left of each bar. (c) shows the predicted protein structures for reverse transcriptases from *Faecalibacterium prausnitzii* infecting phage Taranis (GenBank accession: NC_047914) in blue, *Mycena chlorophos* (AFDB accession AF-A0A146HGT5-F1-v4, sequence identity 19.5%, TM-score 0.87) in yellow and from an uncultured archaeon (AFDB accession AF-A0A1S6HH33-F1-v4, sequence identity 23%, TM-score 0.93) in purple. The structures were trimmed keeping only the residues of the Foldseek alignments and the top hit to a bacterium (AF-A0A2A7AF35-F1-v4) was omitted for clarity as it was near identical (TM-score 0.98). (d) shows the predicted integrase of *Mycobacterium* phage Highbury (GenBank Accession: OR521086) in blue, and predicted recominbases/integrases from *Trichuris trichiura* (AFDB accession AF-A0A077ZEL6-F1-model_v4, sequence identity 21.4%, TM-score 0.92) in yellow and *Enterococcus faecalis* (AFDB accession AF-A0A132P2T2-F1-model_v4, sequence identity 17.9%, TM-score 0.92) in red. The predicted recominbase from *Methanocaldococcus jannaschii* (AFDB accession AF-Q57813-F1-model_v4, sequence identity 22.6%, TM-score 0.78) was omitted for clarity. (e) shows the predicted protein structures for a predicted 2OG-Fe(II) oxygenase on *Synechococcus* phage S-H9-1 (GenBank Accession: NC_070961) in blue and the human Prolyl hydroxylase EGLN3 (UniProt accession: Q9H6Z9) in yellow, with alignment sequence identity 18.6% and TM-score of 0.87. The structures were trimmed keeping only the residues aligned by Foldseek. (f) shows the predicted protein structures for a predicted amidase on *Delftia* phage IME-DE1 (GenBank Accession: NC_028702) in blue and the human Peptidoglycan recognition protein 1 PGLYRP1 (UniProt accession: O75594) in yellow, with alignment sequence identity 16.0% and TM-score of 0.79. The structures were trimmed keeping only the residues aligned by Foldseek.

The Phold DB 3.16M PHROG with the most hits to all three domains was PHROG 1423 (12,673 hits) (Supplementary Tables 15-16), containing diversity generating retroelement reverse transcriptases, unsurprising given the prevalence of retrotransposons and retroviruses in *Eukaryota* and prokaryotes^75^. For example, homologs to the reverse transcriptase present in *Faecalibacterium* phage Taranis (GenBank accession: NC_047914) were found in the agaric fungus *Mycena chlorophos* and an uncultured archaeon with TM-scores of 0.87 and 0.93 despite sequence identities of only 19.5% and 23.0% (Figure 6C).

To focus our analysis on a subset of well-annotated genomes, we repeated the same analysis for the 48 high quality reference proteomes (16 from *Bacteria*, 31 from *Eukayota* and a single archaeon, *Methanocaldococcus jannaschii*) with 564,446 proteins that make up the AlphaFold-Proteome database^76^. Hits to enzymatic and metabolic PHROG categories again dominated, with fewer hits to phage structural categories such as tail and connector compared to hits against AFDB (Supplementary Figure 17). For Phold DB, 345,836 (10.9%) had hits to *Bacteria* proteins, 157,877 (5.0%) to *Eukaryota* and 49,229 (1.6%) to *Methanocaldococcus jannaschii*, with 29,449 (0.9%) having hits to all three. For INPHARED 1419, the numbers of hits were 24,669 (16.7%), 12737 (8.6%) and 3,889 (2.6%) respectively, with 1,913 (1.3%) shared. Of these 1,913, the most common single annotation was ‘integrase’ (212). Many INPHARED 1419 integrases had extremely high (> 0.8) TM-scores to proteins from different domains with sequence identities below 25%, suggesting integrases are extremely conserved structure and function across the tree of life^77^. For instance, the predicted integrase of *Mycobacterium* phage Highbury (GenBank Accession: OR521086) has TM-scores of 0.92, 0.92 and 0.78 with sequence identities of 21.4%, 17.9% and 22.6% to predicted recominbases/integrases from *Trichuris trichiura*, *Methanocaldococcus jannaschii* and *Enterococcus faecalis* respectively (Figure 6D). Other core metabolic enzymes, such as thymidylate synthases, methyltransferases and glycosyltransferases, were commonly found in INPHARED 1419 phages with high structural similarity to proteins from other domains of life (Supplementary Table 16).

Finally, 54,406 Phold DB and 5,552 INPHARED 1419 proteins had *Homo sapiens* hits. Compared to hits shared between all three domains of life, hits to *Homo sapiens* were especially enriched towards phage non-structural PHROG categories of nucleic acid metabolism, other and moron, auxiliary metabolic gene and host takeover proteins (Supplementary Figure 18). The most common PHROG groups from Phold DB 3.16M with *Homo Sapiens* hits generally had core nucleic acid metabolic functions, including to PHROG 1249 acetyltransferase (1423 proteins), PHROG 16 DNA helicase (1,322 proteins) and PHROG 61 (2OG-Fe(II) oxygenase) (Supplementary Tables 17-18).

Analysis of the most hit *Homo Sapiens* protein for both Phold DB (2,459 hits) and INPHARED 1419 (303 hits), Prolyl hydroxylase EGLN3 (UniProt accession Q9H6Z9), revealed widespread distribution of structurally similar proteins throughout phages that infect marine bacteria such as *Synechococcus*, *Vibrio and Cyanobacteria.* These proteins had amino acid sequence identities ranging from 8.4-22.5% with EGLN3, with most annotated as ‘2OG-Fe(II) oxygenase’ by Phold (Figure 6E). Following structural multiple sequence alignment of these hits with FoldMason^78^, we found that the two histidine residues comprising the iron cation binding sites^79^ were highly conserved, while the next most conserved residue was Glutamine, forming part of the conserved 2-His-1-carboxylate facial triad canonical for 2OG-Fe(II) oxygenases^80^. This suggests these proteins are correctly annotated as 2OG-Fe(II) oxygenases, which likely confer evolutionary advantages for these phages adapted to the marine environment. We also found 43 INPHARED 1419 (and 292 Phold DB) proteins with strong structural similarity to the human peptidoglycan recognition protein 1 PGLYRP1 (Uniprot accession O75594). These hits were present across a wide variety of phages annotated as ‘amidase’ or ‘lysin’ by Phold, with 14.5-26.1% amino acid similarity to PGLYRP1. This structural similarity suggests a similar mechanism of peptidoglycan recognition between these phages and PGLYRP1 in *Homo sapiens* (Figure 6F).

Overall, such strong structural similarity between phage proteins encoding for core nucleic acid metabolism and other enzymatic functions, suggests a wealth of folds and functions that are present both in phages and across the tree of life^81^.

## Discussion

The availability of large-scale protein sequence and structure databases, combined with improvements in structural alignment algorithms have allowed for the illumination of difficult-to-annotate protein dark matter^52,82,83^. Phage proteins are amongst the most difficult; sequence-based homology approaches leave over 65% of proteins on an average phage unannotated.

In this study, we present Phold, a framework that combines rapid structural information inference using the ProstT5 pLM with Foldseek, leveraging our database of over 1.36M phage protein structures to provide interpretable structure-based annotation with curated high-quality functional annotation labels. We show that Phold has wide applications to prokaryotic virus annotation, including identification of potential antimicrobial resistance proteins and anti-phage defence proteins. Our structural similarity analyses of Phold DB and annotated cultured phages show that many phage proteins, particularly those with nucleic acid metabolism and other enzymatic functions, have structural homologs across the tree of life, including in *Homo sapiens*. Given Phold’s improved performance over sequence-based homology tools, we anticipate Phold will be extremely valuable for phage, archaeal virus and viral metagenomic research communities. We also believe the functional annotations generated by Phold symbiotically provide a platform for guiding wet-lab studies to confirm or disprove these hypotheses, which can be used to continually improve functional labels for future phage annotation databases.

## Methods

All benchmarking in this manuscript was conducted on a single high-performance computing (HPC) node with a single NVIDIA A100-40G GPU along with 32 threads CPU and 128GB RAM of 2x Intel Xeon Platinum 3460Y CPU system with 36 cores @ 2.4GHz. ProstT5 was run with GPU unless otherwise specified.

### Phold Database Construction

#### PHROG

The core of the Phold database is the Prokaryotic Virus Remote Homologous Groups (PHROG) database^22^. All 868,340 PHROG proteins from the core 38,880 PHROGs containing at least 2 proteins were downloaded and deduplicated to retain 440,550 non-redundant proteins. All proteins over 3000 amino acids (AA) in length were then split into equal sizes below 3,000AA (e.g. a 5000AA protein was split into two 2,500AA parts, an 8,100AA protein was split into three 2,700AA parts) due to this being approximately the memory limit of our available GPUs for structure prediction (NVIDIA A100 40GB on the University of Adelaide Phoenix HPC and AMD MI250x on Setonix at the Pawsey Supercomputing Research Centre), yielding 441,177 unique protein sequences (including fragments) overall. Structure prediction was then conducted for all proteins using two methods. The first was using ColabFold v1.5.3^30^ implementing AlphaFold2^25^, conducted on Phoenix. Specifically, MSAs were created using both the uniref2302_30 and colabfold_envdb_202108_db databases using MMseqs2^11^ v71dd32, and AlphaFold2 was run in batch mode with 3 models (‘--num-models 3’) and the default 3 recycles, without AMBER relaxation. The model with the highest pLDDT was chosen as the best ColabFold model.

Additionally, structures for all PHROG proteins that fit in memory (less than approximately 900AA) were then predicted using ESMFold^31^. The structure prediction with the highest pLDDT between ESMFold and ColabFold models was then selected as the best model to be included in the Phold DB 3.16M.

#### EFAM

To diversify existing PHROG groups, we downloaded all 402,958 extremely conservative efam viral proteins taken from the Global Ocean Virome 2.0 dataset^45^. 392,139 proteins remained after filtering out all proteins with unclassified amino acids, of which 392,079 were lower than 3000AA and kept for inclusion in the Phold DB 3.16M. Protein structure prediction was then conducted for these using ColabFold and ESMFold in an identical fashion as the PHROGs.

To assign efam proteins to a PHROG, the top ranking structure for each efam protein was then compared against the PHROG structures in an all-vs-all search using Foldseek v 6cfb880^26^ using the parameters “-c 0.7 --num-iterations 3” to retain only hits with a bidirectional coverage of 70% (i.e. across the majority of both the query efam and target PHROG protein). All hits were then filtered using a minimum alignment TM-score value (computed by Foldseek) of 0.6 indicating that both the query and target are very likely to share the same fold^73^. For alignments under 75AA, a Foldseek E-value cutoff of 0.01 was also implemented along with the TM-score filter to attempt to avoid spurious protein structure prediction alignments^74^. The top hit PHROG was then assigned for each of the 233,181 efam proteins that passed these criteria, which were included in the Phold Search DB 1.36M. The remaining 158,898 proteins without PHROG hits were not included in the Phold Search DB 1.36M but remain available in the overall Phold DB 3.16M.

#### ENVHOG

To further diversify existing PHROG groups and provide a more general representation of the viral protein space, we downloaded MMseqs2 position-specific scoring matrix (PSSM) profiles for all of the Environmental Viral Homologous Groups (enVhogs) accessed on 31 January 2025 from http://envhog.u-ga.fr/envhog/^44^. Consensus sequences for each profile were extracted for each of the 2,203,457 enVhogs using the “mmseqs profile2consensus” command using MMSeqs2 v16-747e6^11^. All enVhogs over 3000AA were fragmented in the same fashion as PHROGs before structure prediction, yielding 2,205,969 sequences overall.

Protein structure predictions were then generated for each consensus enVhog using the same methodology as for PHROGs using ColabFold and ESMFold, with two differences. The first difference is that Colabfold v1.5.5 was used with the AMD MI250x GPUs on Setonix in a Singularity container available from https://quay.io/repository/sarahbeecroft9/colabfold/rocm6.0.0_cpuTF, but otherwise identical parameters were used for protein structure generation. The second difference was that MSA generation was done with MMseqs2 v15.6f452^11^ with a third database added to enrich the MSA generation pipeline using beyond ColabFold’s default uniref2302_30 and colabfold_envdb_202108_db databases to generate deeper MSAs for these proteins.

Implementation to search using this third “colabfoldv” database is available at https://github.com/gbouras13/colabfoldv and contains 129944764 non-redundant viral (predominantly phage) proteins. Structure prediction was also conducted for these using ESMFold, with the top-ranking protein chosen in an identical fashion as for the PHROGs.

Assignment of enVhog proteins to PHROG annotations was done using two methods. Firstly, the top ranking predicted structure for each enVhog protein was then compared against the PHROG structures in an all-vs-all search using Foldseek v6cfb88^26^ using the parameters “-c 0.7 --num-iterations 3” to retain only hits with a bidirectional coverage of 70% (i.e. across the majority of both the query enVhog and target PHROG protein). All hits were then filtered with a minimum alignment TM-score value computed by Foldseek of 0.6 indicating that both the query and target are very likely to share the same fold^73^. For alignments under 75AA, a Foldseek E-value cutoff of 0.01 was also implemented to avoid spurious protein structure prediction alignments as for efam^45^. Secondly, a HMM-HMM comparison approach using HHblits (via hhsuite v3.3.0)^84^ implemented using the same methods as in the enVhog manuscript was conducted using an E-value of 0.001. However, to increase the confidence in the functional PHROG assignment by considering coverage across the entire protein, we added a filter to HMM-HMM hits, ensuring at least 70% coverage of both the query HMM and target HMM consensus sequences.

In the case that an enVhog had both Foldseek and HMM-HMM hits to different PHROGs, the hit with lowest E-value was taken as the best hit for that enVhog. Overall, 562,369 enVhog consensus proteins with either Foldseek or HMM-HMM hits to PHROG groups with known function were assigned. Of these, 383,648 were unique to Foldseek, 48,570 unique to HMM-HMM comparions, and 130,151 had hits with both methods. Of these 130,151 with hits to both methods, 94,526 had hits to identical PHROGs, 9,951 had hits to different PHROGs with identical annotations, 24,035 had hits to different PHROGs with the same PHROG category but different specific annotations, 8,527 had hits to different PHROG categories where one hit was to an unknown function PHROG category (in which case the known function hit was taken), while 3,063 had conflicting hits to different PHROG categories where both had known annotations. The remaining 1,638,667 enVhog proteins without PHROG hits were not included in the Phold Search DB 1.36M but remain available in the overall Phold DB 3.16M.

#### PHROG SINGLETONS

To improve the functional understanding of singleton PHROGs, we took all PHROG singleton proteins (i.e. from PHROG 38,881 to 109,404) and initially generated predicted protein structures using ColabFold and ESMFold in the same way as described for PHROGs above. Due to the likelihood for these singletons to have sparser MSAs and lower predicted pLDDT, we also generated more predicted structures using enriched MSAs to increase the chances of generating a high confidence structure. Specifically, we generated MSAs using both the standard ColabFold databases (i.e. uniref2302_30 and colabfold_envdb_202108_db) and the standard ColabFold databases augmented with the “colabfoldv” database in the same way as for the enVhogs using MMseqs2 v15.6f452^11^. The protein structure with the highest pLDDT between the six ColabFold models (three with default MSA, three with enriched MSA) and the ESMFold model was chosen as the best structure.

#### Anti-CRISPRdb

All 3693 Alphafold2 predictions from Sahakyan et al^85^ of anti-CRISPRdb proteins were downloaded from https://zenodo.org/records/7747008. 3652 predicted anti-CRISPR structures with average pLDDT over 70 (from the metadata as provided by Sahakyan et al) were kept for inclusion in Phold. Predicted structures for these were also generated with ESMfold. The protein structure with the highest average pLDDT was then included in Phold’s databases.

#### CARD

The CARD Data v3.2.8 release (October 2 2023) was downloaded from https://card.mcmaster.ca/download^47^. 4,804 protein homolog models were considered for inclusion in Phold. 2,893 of these proteins had structure predictions already in the AlphaFold database v4^51^, which were downloaded. Structure predictions for the remaining 1,911 proteins were generated on Phoenix in the same way as the PHROGs. Predicted structures for these were also generated with ESMFold. The protein structure with the highest average pLDDT was then included in Phold’s databases.

#### DefenseFinder

All 461 monomer DefenseFinder^48^ protein structures generated using Alphafold2 available as of January 31 2024 were downloaded from https://defensefinder.mdmlab.fr/. 408 of these with average pLDDT above 70 were kept. Predicted structures for these were also generated with ESMfold. The protein structure with the highest average pLDDT was then included in Phold’s databases.

#### Diversity-Generating Retroelement (DGRs) Reverse Transcriptases

To improve the diversity of DGRs in Phold’s databases, we took 12760 putative DGR reverse transcriptase sequences from Roux et al^46^ representing all OTUs (≥95% AAI) that corresponded to DGR references. We generated predicted protein structures for all 12760 in the same way as for PHROGs on Phoenix. We then ran Foldseek easy-search v9.427df8a^26^ to compare the best predicted structure based on pLDDT against all PHROG 1423 (annotated as reverse transcriptase) using ColabFold predicted protein structures with an E-value threshold of 1E-05. 12,683 out of 12,670 putative DGRs with hits were then kept for inclusion in Phold. Predicted structures for these were also generated with ESMFold. Finally, the protein structure with the highest average pLDDT was then included in Phold’s database.

#### NetFlax

We used the NetFlax^50^ toxin-antitoxin proteins to improve the diversity of toxin-antitoxin system proteins in the Phold database. We took all 7,152 toxin and antitoxin protein structures predicted with AlphaFold2 from NetFlax as provided by the authors. Predicted structures for these were also generated with ESMFold. Finally, the protein structure with the highest average pLDDT was then included in Phold’s databases.

#### Virulence Factor Database

We generated protein structure predictions for the Virulence Factor Database (VFDB)^49^ (downloaded on 19 January 2024 from http://www.mgc.ac.cn/VFs/download.htm) for 27,823 VFDB virulence factor proteins using the same methods as PHROGs. The protein structure with the highest average pLDDT between ColabFold and ESMFold methods was then included in Phold’s databases.

### Phold Benchmarking Datasets

All per genome and per protein structure information for the benchmarking datasets can be found in Supplementary Tables 14, 19 and 20.

We used 5 different datasets for benchmarking Phold:

1. 182 INPHARED^10^ viral genomes with a total of 16460 CDS (‘INPHARD 182’).
Specifically, these were comprised of one representative of all unique genera added to INPHARED after the 1 October 2021 release as of the 1 November 2023 release. We chose 1 October 2021 as a timepoint cutoff, as theoretically phages added after 1 October 2021 were not part of the original PHROGs database, though we note that we do not guarantee this, due to the frequent taxonomic renaming of phages and constant updating of INPHARED. We used this dataset to broadly capture the known diversity of reasonably well-characterised, mostly isolated phages and archaeal viruses in place of the 1419 INPHARED dataset for all Phold performance ablations, due to its smaller size.
2. 1419 INPHARED viral genomes comprising one representative for every unique viral genus in INPHARED as of the 14 April 2025 release, containing 148,196 CDS (‘INPHARED 1419’).
3. <u>249</u> *Crassvirales* phage genomes from Edwards et al^6^ with a total of 22,127 CDS (‘Crass’).
4. 63 metagenomic assembled phage genomes from Cook et al^3^ with 3,699 CDS (‘Cook’).
5. 45 metagenomic assembled phage genomes from Tara Oceans^59^ with 2,112 CDS (‘Tara’).

For each benchmarking dataset, all phage genomes were first run with Pharokka v1.7.4 or v1.7.5^16^ using pyrodigal-gv^18,56,57^ as the genecaller with ‘-m’ (meta mode) with and without ‘--meta_hmm’ that implements PyHMMER^15^ based annotation in addition to MMSeqs2 profile-based annotation. Protein structure predictions for all predicted coding sequences (CDS) were folded using ColabFold v1.5.3 and ESMFold using the same parameters as the PHROGs for database construction, though in this case, the top ranking pLDDT model was not chosen, as ESMFold and ColabFold were considered differently in benchmarking (i.e. the ColabFold model was used as the pseudo ground truth). A small number of large CDS over 3000AA in each dataset due to computational limitations and these were ignored in benchmarking. Phold parameters ‘--structures --structure_dir’ with ColabFold-generated structures, along with the default ‘--max_seqs 1000 –masking_threshold 25’ were used with the Phold Search DB 1.36M and an E-value threshold of 0.01 were used to generate the baseline pseudo ‘ground-truth’ for all sensitivity analyses. For the purposes of benchmarking, proteins with ground truth non-hypothetical functional annotations were considered as true positives. Of the metrics presented, the binary task indicates whether the 3Di inference method could generate any non-hypothetical annotation for the proteins, while multiclass metrics indicate that the method generated the correct specific annotation (i.e. at either PHROG group, product or category level).

More information on each of the metagenomic phage datasets are as follows:

#### *Crassvirales* Phages (‘Crass’)

249 *Crassvirales* genomes were taken from Edwards et al^6^. Protein structure predictionx were generated for all proteins 2000 amino acids and below, with a total of 21,740 protein structure predictions were generated.

#### Cook et al Phages ‘(Cook’)

All putative phage genomes assembled with Phables v0.2.0^86^ and MEGAHIT v1.2.9^87^ in the file BINS_sample_01_phables.fa.gz from Cook et al were downloaded. CheckV v1.0.3^88^ was run to assess the completeness of these genomes. 64 genomes were assessed to be complete. One genome (phage_comp_52_cycle_2) was excluded as being very similar to another genome (phage_comp_52_cycle_1) based on the Phables assembly method from the manuscript. Of 63 remaining genomes, predicted structures were generated for 3,695 out of 3,699 proteins.

#### Tara Oceans Phages (‘Tara’)

Reads from Sequence Read Archive (SRA) accessions ERR2750826, ERR2750828 and ERR2750829 were downloaded and assembled with Phables^84^ v1.3.0, and metaSPAdes v3.15.5^89^ yielding 45 phage genomes. Predicted structures were generated for 2,110 out of 2,112 predicted phage genomes.

### Phold Search Parameter Optimisation and Comparison with Foldseek-GPU

To understand the effect of differing Foldseek^26^ search parameters on search sensitivity and resource consumption, we ablated the ‘--max-seqs’ parameter (controlling search sensitivity), choosing 1000 (default in Foldseek), 5000, 10000, 20000, 50000 and 250000. We chose to do this using both Foldseek CPU and GPU, as it is likely that the many Phold users would use either implementation depending on their hardware. We also clustered the Phold search database at different values of ‘--seq-id’ (0,0.1,0.2,0.25,0.3,0.4,0.5,0.6,0.7,0.8,0.9) and searched the clustered database using the ‘--cluster-search 1’ parameters and compared it to a full ‘Foldseek search’ of the entire Phold search database. We performed the same ablations against both the Phold Search DB 1.36M (i.e excluding enVhogs and efam proteins with no PHROG hits) consisting of 1,363,704 proteins and the full Phold DB 3.16M (i.e. including efam and enVhog proteins with no PHROG hits) consisting of 3,166,202 proteins. The query dataset used for this was INPHARED 182. Phold with ProstT5 with the commit ‘59d356è (with customised clustered databases as described above) was used. For Foldseek GPU searches, we specified the Phold parameter ‘--foldseek_gpù, which passes ‘--gpu 1 -- prefilter-mode 1’ to Foldseek. The metrics compared were total wall-clock time taken, total CDS with functional hits found (i.e. hits to proteins with a function that was not unknown function) and total CDS with all hits (i.e. including hits to proteins with unknown function). Foldseek v10.941cd33 with an E-value of 0.001 was used for Phold runs.

### Phold ProstT5 Confidence Masking Benchmarking

We define the per-residue ProstT5 confidence as a score out of 100. It is calculated as 100 multiplied by the softmax of the logits resulting from the ProstT5-CNN head for the 3Di token with the highest value that is chosen at that residue. The per-protein ProstT5 confidence metric is accordingly defined as the mean of each per-reside confidence score across the protein. It reflects a measure of the model’s confidence in the 3Di prediction at each residue. In order to understand the effect of masking low confidence ProstT5 predicted residues on Phold’s search sensitivity, we conducted an ablation study with the ‘--mask_threshold’ parameter. The ‘--mask_threshold’ masks 3Di tokens in the Foldseek search for residues where the ProstT5 confidence is below this threshold. We tested values of 0 (i.e. no masking), 10, 20, 30, 40, 50, 60, 70 and 80 and compared the number of CDS annotated compared to running Phold with the ColabFold generated structures against the search database, using an E-value of 0.001 and max-seqs 1000.

### ProstT5 Confidence Analysis

For the 3,166,202 protein structure predictions in Phold DB 3.16M, we ran ‘phold predict’ with ProstT5-CNN to predict 3Di tokens. For each protein, the mean ProstT5 3Di confidence, calculated by Phold as the mean of all per-residue ProstT5 confidence scores across the protein was taken. The actual 3Di similarity was calculated by computing the edit distance between ProstT5 generated 3Di and the 3Di tokens generated by the Foldseek VQ-VAE from the 3,166,202 protein structure predictions in Phold DB 3.16M. Pearson’s r and Spearman ρ were then calculated for between the ProstT5 confidence and actual similarities.

### ProstT5 LoRA Finetuning and CNN Training

To finetune ProstT5, 2,205,504 predicted protein structures with a mean pLDDT of at least 70 were taken from Phold DB 3.16M protein structures. They were clustered with Foldseek v10.941cd33 using the ‘Foldseek cluster’ command using the default bi-directional coverage of 80%, yielding 149,321 non-singleton and 625,617 singleton clusters. To avoid oversampling large clusters in the training dataset, only the 20 most diverse sequences for non-singleton clusters with at least 20 members were chosen. For training, proteins larger than 512 amino acids were truncated to fit in GPU VRAM, keeping only the first 512 residues. We held-out 100 random non-singleton clusters with 719 proteins and 625 random singletons as the validation set, with the remaining 1,323,937 proteins comprising the training set. We also used 500 arbitrary Swiss-Prot proteins as a non-phage validation set. We applied low order rank adaptation (LoRA) fine-tuning^90^ on the full ProstT5 encoder-decoder with weights accessed via HuggingFace. Specifically, our configuration was the same as suggested by Schmirler et al^91^ : rank 4, alpha 1, applied to query, key, value, and output of the attention layers. Overall, out of 2,818,852,864 (2.8B) ProstT5 encoder-decoder parameters, 5,900,288 (5.9M) were available for fine-tuning. Our fine-tuned model was trained with Pytorch v2.5.1 using the default HuggingFace trainer on a single GPU node containing 4x NVIDIA A100 40GB GPUs. We trained our model for 4 epochs with an effective batch size of 32 (per GPU minibatch size of 1, gradient accumulation steps of 8), using mixed precision, a learning rate of 3e-4 and no warmup. The fine-tuned model is freely available at https://huggingface.co/gbouras13/ProstT5Phold.

We then trained two-layer convolutional neural networks (CNNs) on top of the LoRA finetuned ProstT5 encoder embeddings using biotrainer^41^. We used identical hyperparameters to the CNN trained and provided with the original ProstT5 (two layers, bottleneck dimension 32, no drop out, batch size 8, 5 epochs and a learning rate of 1e-3) and trained the CNNs on a single NVIDIA RTX4090 GPU. We trained two CNNs: one using the “CASP14” dataset from ProstT5 containing proteins from the protein data bank (PDB) along with some CASP14 proteins named ‘vanilla CNN’ in this manuscript; and one using 50,000 randomly selected Phold DB 3.16M proteins taken from the 1,323,937 proteins in the LoRA finetuning training dataset named ‘finetune CNN’ in this manuscript. These CNNs are available in the Phold GitHub repository. We ultimately chose to keep the original ProstT5 with CNN for use with Phold (see Supplementary Note 2).

### Phold DB Database Curation and Unknown Function PHROG Annotation Propagation

Original PHROGS with Two or More Members (1-38880)

For all PHROG proteins that had at least two members in the original PHROGs v4 (i.e. from PHROG 1 to 38880) which were annotated as “unknown function”, a semi-manual curation approach was as follows. Overall, 668,199 proteins (i.e. original PHROGs expanded with efam and enVhog proteins) belonged to the 33,553 PHROGs that were “unknown function” in v4, while 651,666 proteins belonged to the 5,327 PHROGs that had an annotated function.

We ran Foldseek (CPU) easy-search with the parameters ‘-c 0.7 --max-seqs 1000 --num-iterations 3’ against the following 3 databases:

1. All BFVD structures.
2. All AFDB structures (searched using the Foldseek AFDB50 database with --cluster-search 1)
3. All 651,666 proteins belong to the 5,327 known function PHROGs.

We then post filtered all the Foldseek results to keep all hits that had E-value less than 0.01 and a TM-score of at least 0.6, indicating that the query and target likely shared the same fold^73^. Annotation transfer for PHROGs that were annotated as “unknown function” in v4 was then conducted manually using the following heuristics:

- For each PHROG group, number and strength (in terms of E-value TM-score and LDDT) of the top hit target against 1) the BFVD and 2) the known PHROGs were considered as the best source of information, as these databases most tailored for viral and phage proteins. For any annotation transfer to take place, at a minimum, at least 10% of query proteins in the PHROG must have had an identical or very similar annotation to considered (though generally a higher amount was needed to transfer annotation)
- Generally, vague annotations (e.g. those with ‘DUF’, ‘Uncharacterized’, ‘Hypothetical’, ‘Putative’) and hits near E-value (0.01), TM-score (0.6) and percentage (10%) thresholds were not propagated. Vague annotations (e.g. uncharacterized) were generally not counted as informative (i.e. not penalised as divergent) if the rest of the annotations were informative and consistent.
- PHROGs with queries that had multiple divergent annotations were not propagated.
- Other PHROG level information computed in the original PHROG manuscript and available for download and on the webserver was also considered as informative. These particularly include the annotation of similar PHROGs computed using HMM-HMM searches and similar PFAM, KEGG and GO hits.
- If a PHROG had only 1 or 2 members, then a strict 100% criterion was enforced and if 3 members, a 2/3 criterion was applied.
- Putative integrases were specifically scrutinised in more detail including visualising and comparing hits manually, given the therapeutic interest in identifying them.
- BFVD and similar PHROG information was supplemented by AFDB top hit counts and information. This was particularly used for PHROG queries that had no hits to BFVD proteins, and for PHROGs likely to belong to non-structural categories (i.e. other, moron, auxiliary metabolic gene and host takeover, DNA, RNA and nucleotide metabolism and transposases).

#### Original Singleton PHROGS (38881-109399)

For PHROG singleton proteins (i.e. PHROG 38881 to 109399), the semi-manual functional curation approach followed the same Foldseek easy-search parameters as for PHROGs 1-38880. The heuristics followed for annotation transfer for these PHROGs were as follows:

- Consider the number and strength of hits of the annotation against the PHROGs 1-38880 as the post important datapoint. Singletons with only few hits and hits near the E-value and TM-score thresholds did not have their annotation propagated.
- Annotation transfer was also not conducted if there were many hits for the same singleton with divergent annotations.
- Also consider the AFDB and BFVD top hit annotation for each singleton, if they exist. If they are strongly divergent, then do not proceed with annotation transfer.
- Manually view the predicted structure to see if it visually concords with the predicted function.

### Low, Medium and High Phold Annotation Confidence Heuristics

To make the output of Phold more interpretable for users, particularly those without understanding of protein structural alignment methods, we assign each CDS annotation as either ‘high’, ‘medium’ or ‘low’ confidence based on the following heuristics. High confidence hits are where both the query and target proteins have at least 80% reciprocal alignment coverage, along with either (i) greater than 30% amino acid sequence identity (suggesting the hit is in the light zone of sequence homology^27^) or (ii) query mean ProstT5 confidence of at least 60% (suggesting a very good quality ProstT5 3Di prediction) or (iii) an alignment E-value < 1e-10. Medium confidence hits are where either the query or target protein hit has at least 80% coverage along with either (i) greater than 30% amino acid sequence identity or (ii) ProstT5 confidence between 45% and 60% (suggesting a good quality ProstT5 3Di prediction) and (iii) an E-value < 1-e05. Low confidence hits are all other hits below the specified E-value that do not fit those thresholds (i.e. hits with low coverage, low amino acid sequence identity, low ProstT5 confidence, or hits near the E-value threshold of 0.001). If Phold is run with user provided input structures instead of ProstT5, the heuristics are identical other than the ProstT5 confidence criteria. In this case, Phold will also output alignment template modeling score (TM-score)^92^ and local distance difference test (LDDT)^93^ values from Foldseek, which may also guide the user in assessing annotation quality.

### NEFF and MSA Analysis

The number of homologs was calculated as the number of MMSeqs2 hits found by ColabFold against both the Uniref50 and environmental databases. The NEFF for each MSA was calculated using NEFFy^94^.

### Phold DB and INPHARED Tree of Life Homology Analyses

To compare protein structures across the tree of life, we took all Phold DB 3.16M and 147,946 INPHARED protein structures and ran Foldseek (CPU) easy-search with the parameters ‘-c 0.7 --max-seqs 1000 --num-iterations 3’ against the following 2 databases:

1. AlphaFold Database^51^ with over 214 million proteins (searched using the Foldseek AFDB50 database with --cluster-search 1).
2. AlphaFold Proteome database consisting of 48 high quality reference proteomes (16 from *Bacteria*, 31 from *Eukayota* and a single archaeon, *Methanocaldococcus jannaschii*) with 564,446 proteins^76^.

We then post-filtered all the Foldseek results to keep all hits that had E-value less than 0.01 and a TM-score of at least 0.6, indicating that the query and target likely shared the same fold.^70^ Taxonomic labels for all hits were taken from the Foldseek output. *Homo sapiens* hits were post-filtered and taken from the Proteome searches, as the smaller target database size led to substantially more hits to the specific taxon (i.e. 17,041 Phold DB proteins had *Homo Sapiens* hits when post-filtering AFDB results compared to 54,406 when post-filtering the Proteome search). We also kept the Foldseek-computed local distance difference test (LDDT) score for every alignment.

## Data Availability

Protein structure predictions for all Phold database proteins can be found at https://doi.org/10.5281/zenodo.16739199 (all Phold Search DB 1.36M structures) and https://doi.org/10.5281/zenodo.16741650 (all additional efam and enVhog structures without PHROG assignment in Phold DB 3.16M but not Phold Search DB 1.36M). Protein structure predictions, genomes and GenBank formatted annotation files for INPHARED 1419 viruses and the Cook, Crass and Tara benchmarking datasets can be found at https://doi.org/10.5281/zenodo.16744439. Foldseek formatted Phold databases are available at https://doi.org/10.5281/zenodo.16741548 and can be downloaded using ‘phold install’.

## Code Availability

Phold is open-source software available at https://github.com/gbouras13/phold. All other code required to recreate the results in this manuscript can be found at https://github.com/gbouras13/phold-analysis. A modified version of the ColabFold MSA search code using an additional database containing 129944764 non-redundant viral (predominantly phage) proteins is available at https://github.com/gbouras13/colabfoldv, while the database can be downloaded from https://doi.org/10.5281/zenodo.15045387.

## Acknowledgements

This work was supported with the assistance of resources and services from Phoenix HPC at the University of Adelaide and Pawsey Supercomputing Research Centre, which is supported by the Australian Government. We would like to thank Fabien Voisin and Sarah Beecroft for their assistance in operating ColabFold at scale at Phoenix and Pawsey respectively, with extra acknowledgement to Sarah for containerising ColabFold for use on Setonix’s AMD GPUs. We would also like to thank Gemma Atkinson and Artyom Egorov for providing us with NetFlax protein structure predictions. R.A.E was supported by awards from the Australian Research Council DP250103825 and FL250100019. M.S. acknowledges support by the National Research Foundation of Korea (grants 2020M3A9G7103933, RS-2021-NR061659 and RS-2021-NR056571 and RS-2024-00396026), Samsung DS Research Fund, Creative-Pioneering Researchers Program, AI-Bio Research Grant through Seoul National University, and Novo Nordisk Foundation (NNF24SA0092560). M.M. acknowledges support from the National Research Foundation of Korea (grant RS-2023-00250470).

