## Supplementary Figures for "Protein Structure Informed Bacteriophage Genome Annotation with Phold"

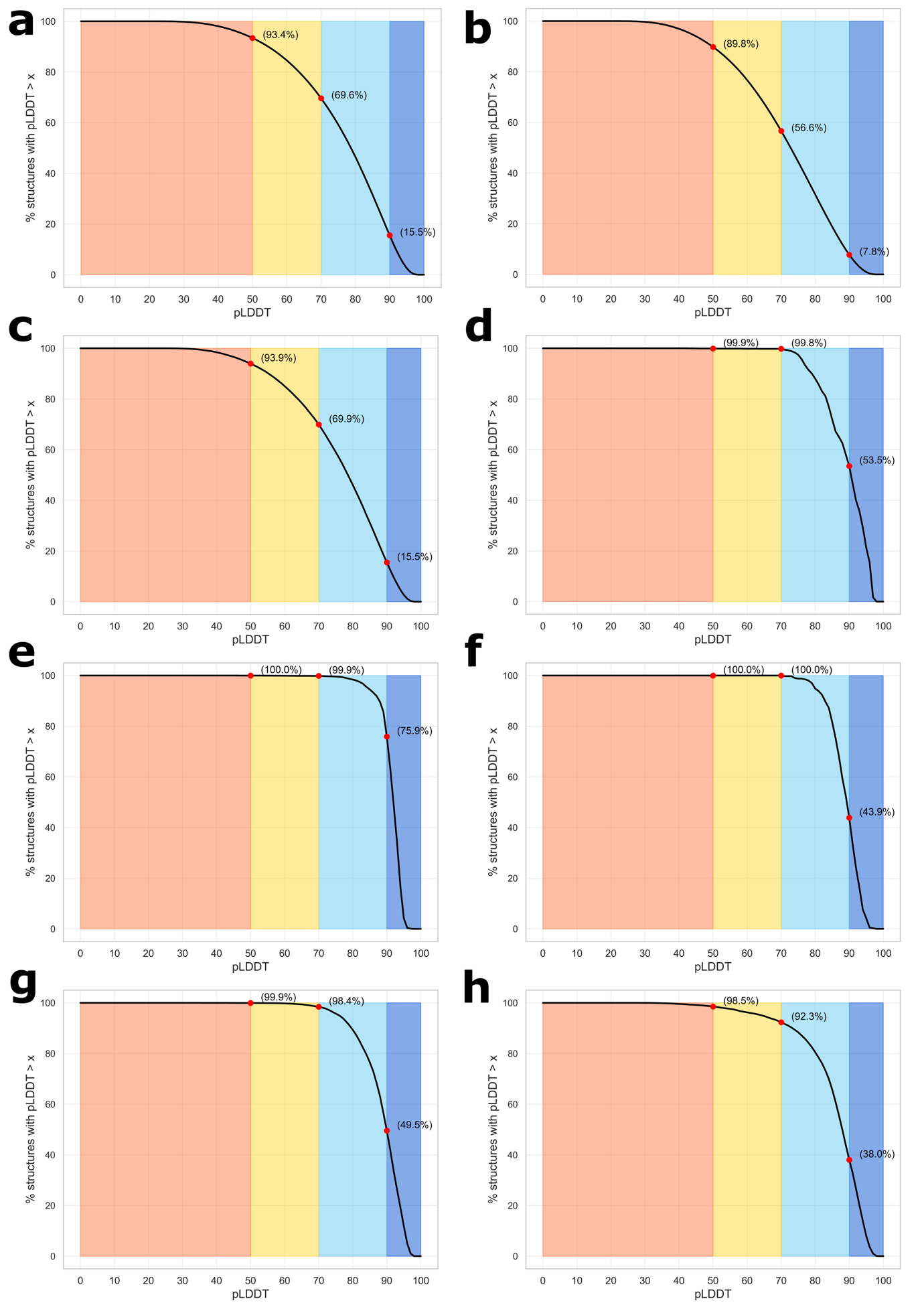

Supplementary Figure 1

Phold sub-database predicted structure quality. For all plots, the proportion of protein structures is (y-axis) plotted against the pLDDT value (x-axis). Colours follow the AlphaFold2 pLDDT colour scheme as implemented in ColabFold’s visualisation.

(a) Phold Database 3.16M (b) 1.78 million enVhog and efam proteins with no assigned PHROG group (i.e. all proteins in Phold Database 3.16M but not in the Phold Search Database 1.37M) (c) PHROG singletons (d) Anti-CRISPR proteins (ACRs) (e) CARD antimicrobial resistance proteins (f) DefenseFinder proteins (g) Netflax toxin-antitoxin proteins (h) VFDB proteins.

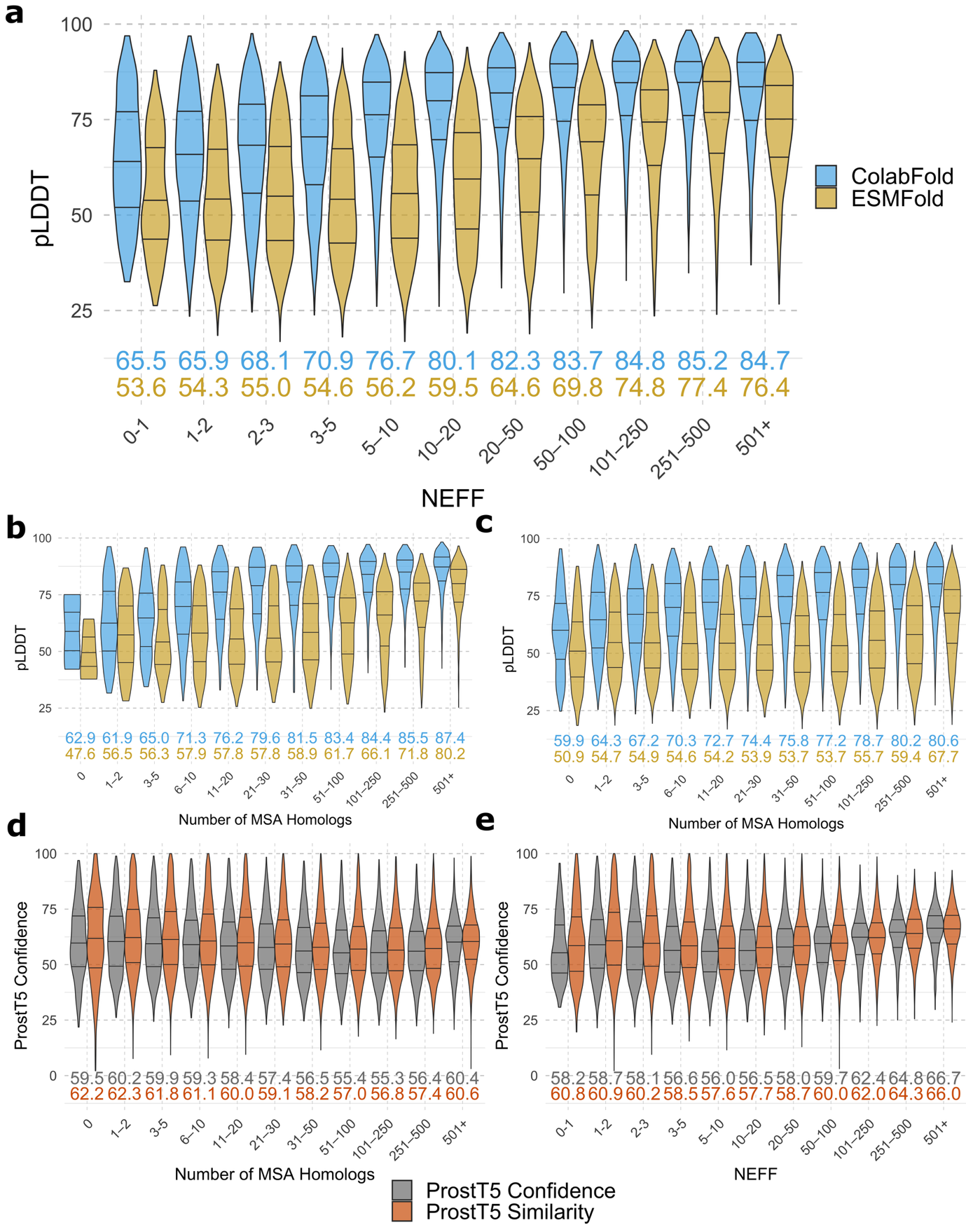

Supplementary Figure 2

Relationship between MSA depth (number of homologs) or quality (NEFF) and various protein structure prediction metrics and datasets in the Phold Database 3.16M. On each plot, medians are displayed numerically coloured by their metric. (a) Violin plots showing pLDDT for ColabFold (blue) and ESMFold (gold) on the y-axis against NEFF (x-axis) bins for all PHROG, enVhog, efam and PHROG singletons proteins (n=3,099,094), (b) number of homologs (x-axis) bins against pLDDT for PHROGs only (n=441,177), (c) number of homologs (x-axis) against pLDDT for enVhogs, efam and PHROG singletons (n= 2,657,917), (d) number of homologs (x-axis) against ProstT5 confidence (grey) and actual 3Di percentage similarity of ProstT5 3Di predictions to ColabFold 3Di (orange) for all PHROG, enVhog, efam and PHROG singletons proteins (n=3,099,094), (e) Same as (d) but with NEFF on the x-axis.

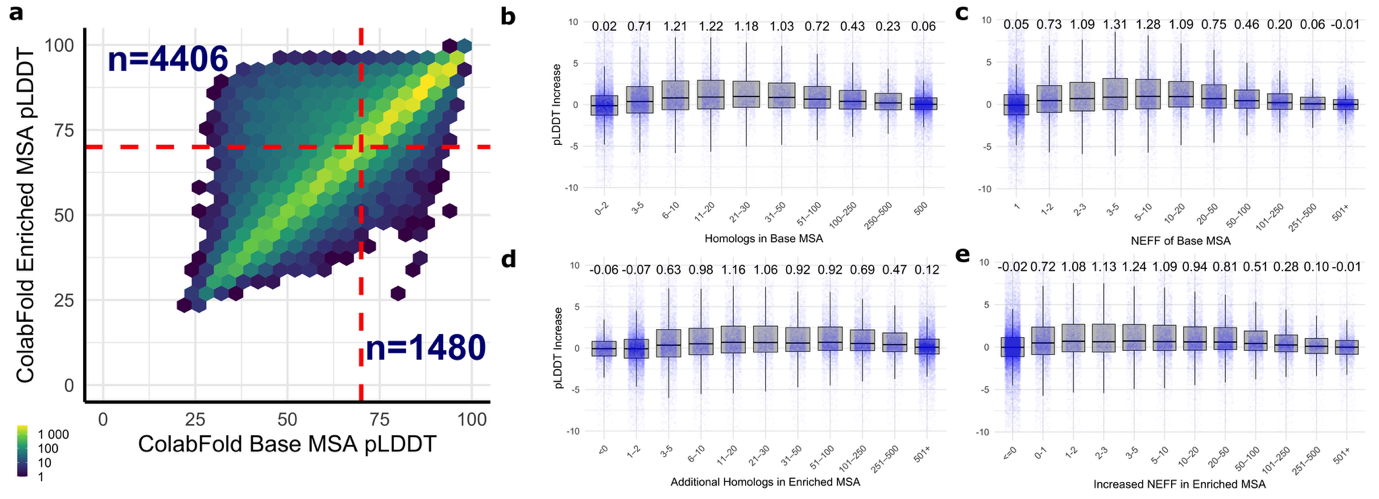

Supplementary Figure 3

PHROG singleton base MSA and enriched MSA analysis. (a) shows a hexbin plot showing the ColabFold structure prediction pLDDT frequency for the base MSA (x-axis) and the enriched MSA (y-axis). (b) contains boxplots showing the difference in pLDDT of ColabFold structure predictions based off the enriched MSA compared to the base MSA (y-axis) compared to the number of homologs in the base MSA. (c) is the same as (b) but the x-axis shows the NEFF of the base MSA. (d) shows the difference in pLDDT of ColabFold structure predictions based off the enriched MSA compared to the base MSA (y-axis) compared to the number of additional homologs in the enriched MSA compared to the base (x-axis). (e) is the same as (d) but the x-axis shows the difference in NEFF between enriched and base MSAs.

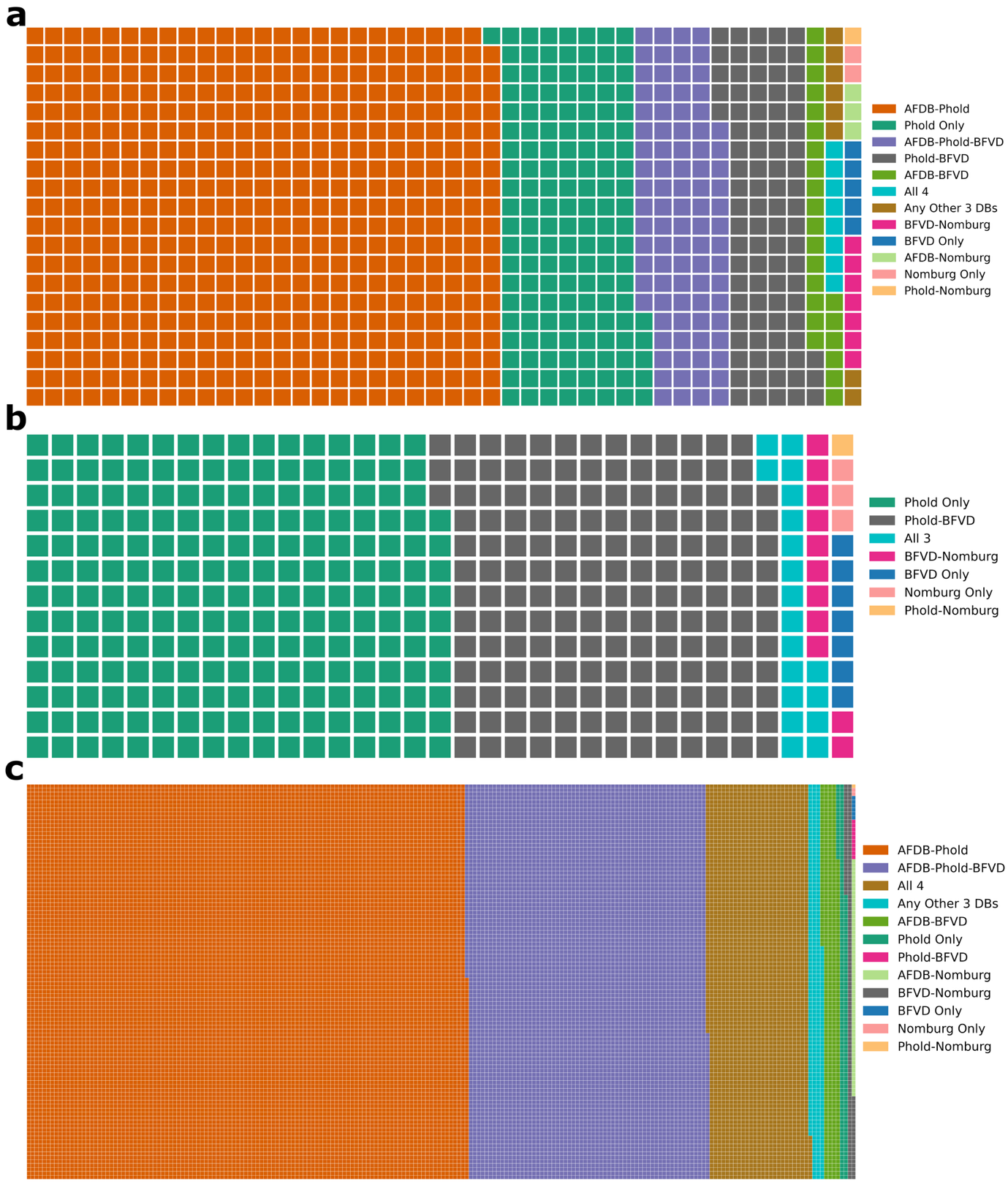

Supplementary Figure 4

Waffle plots. Figure (a) the number of Foldseek clusters containing at least one viral protein shared between AFDB50 representatives, Phold DB 3.16M, BFVD and Nomburg. One square represents approximately 500 clusters. Figure (b) shows the number of viral database-only cluster members containing at least one member shared between Phold DB 3.16M, BFVD and Nomburg. One square represents approximately 1000 proteins. Figure (c) shows the number of cluster members containing at least one member shared between AFDB50 representatives, Phold DB 3.16M, BFVD and Nomburg. One square represents approximately 1125 proteins.

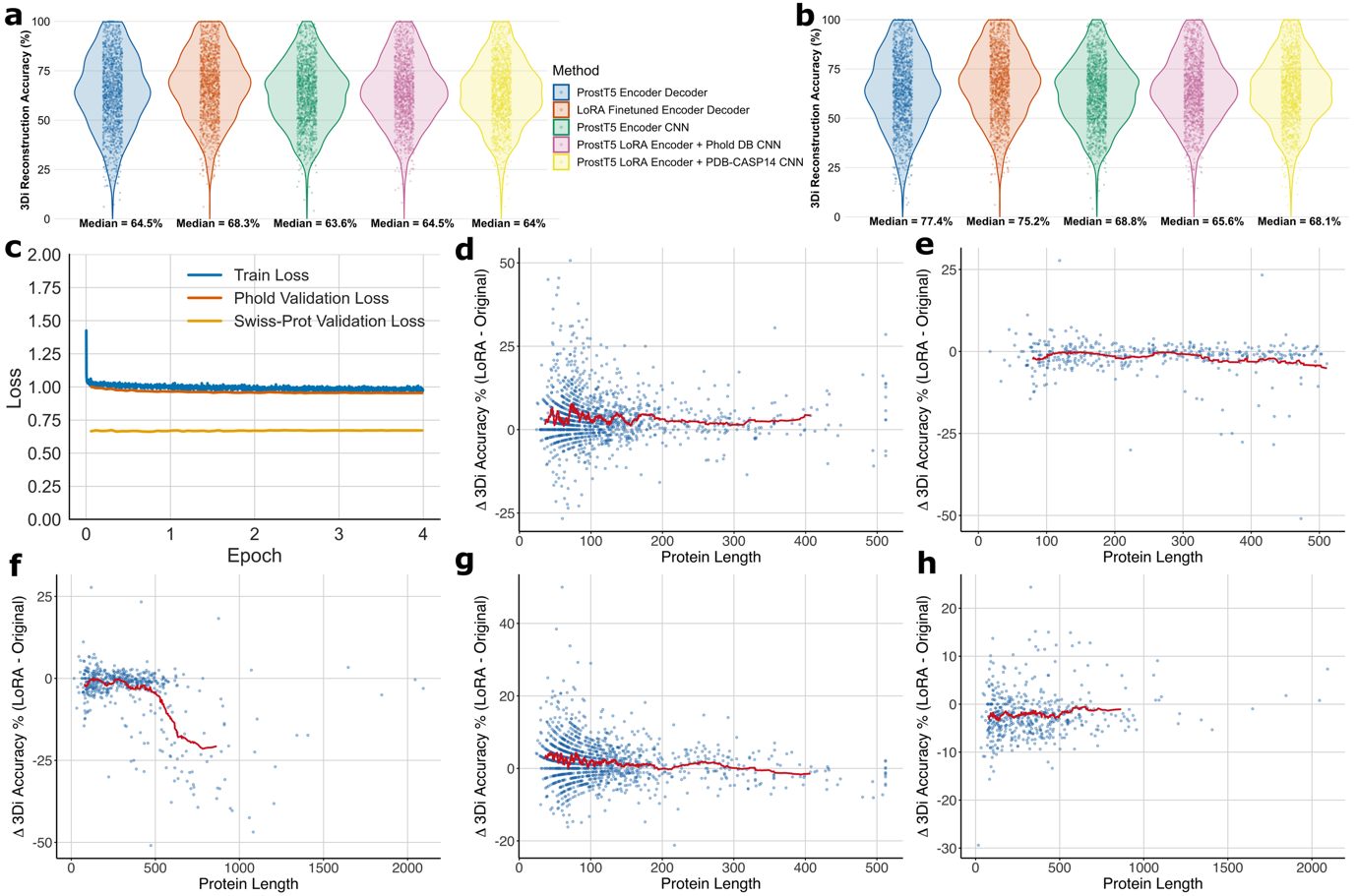

Supplementary Figure 5

LoRA finetuned ProstT5 model benchmarking. (a) and (b) show the 3Di reconstruction performance of LoRA finetuned ProstT5 encoder-decoder and ProstT5 encoder only plus CNN models compared to ground-truth on (a) 1,344 Phold DB validation set proteins and (b) 500 arbitrary Swiss-Prot validation proteins, from left to right: original ProstT5 encoder-decoder (blue); LoRA finetuned ProstT5 encoder-decoder (red); original ProstT5 encoder and CNN (green); LoRA finetuned ProstT5 encoder with CNN trained on Phold DB proteins (purple) and LoRA finetuned ProstT5 encoder with CNN trained on PDB/CASP14 proteins (yellow). Note that generating 3Di tokens for the encoder-decoder models took approximately 53 and 35 minutes wallclock on an NVIDIA A100 40GB GPU for the Phold DB validation set proteins and Swiss-Prot respectively, compared to 36 and 19 seconds for the encoder plus CNN models. (c) shows LoRA finetuned ProstT5 encoder-decoder model losses for training, Phold DB validation and Swiss-Prot validation datasets across the 4 epochs of training. (d)-(h) show the relation between protein length on the x-axis against the difference in 3Di reconstruction accuracy of the LoRA finetuned model compared to the standard ProstT5 on the y-axis, with the red line indicating the last 50-protein moving average) for (d) LoRA ProstT5 encoder-decoder minus the standard ProstT5 encoder-decoder on the Phold DB validation dataset (1344 proteins) (e) LoRA ProstT5 encoder-decoder minus the standard ProstT5 encoder-decoder on the Swiss-Prot validation dataset for proteins 0-512AA long (415 proteins) (f) LoRA ProstT5 encoder-decoder minus the standard ProstT5 encoder-decoder on the Swiss-Prot validation dataset for all proteins (500 proteins), noting that the performance falls dramatically for proteins above the LoRA ProstT5 finetuning dataset crop-size of 512 AA (g) LoRA ProstT5 encoder with CNN trained on Phold Database proteins minus the standard ProstT5 and CNN on the Phold DB validation dataset (1344 proteins) (h) ) LoRA ProstT5 encoder with CNN trained on Phold Database proteins minus the standard ProstT5 and CNN on the Swiss-Prot validation dataset (500 proteins), noting that there is no fall in performance for longer proteins in contrast to (f).

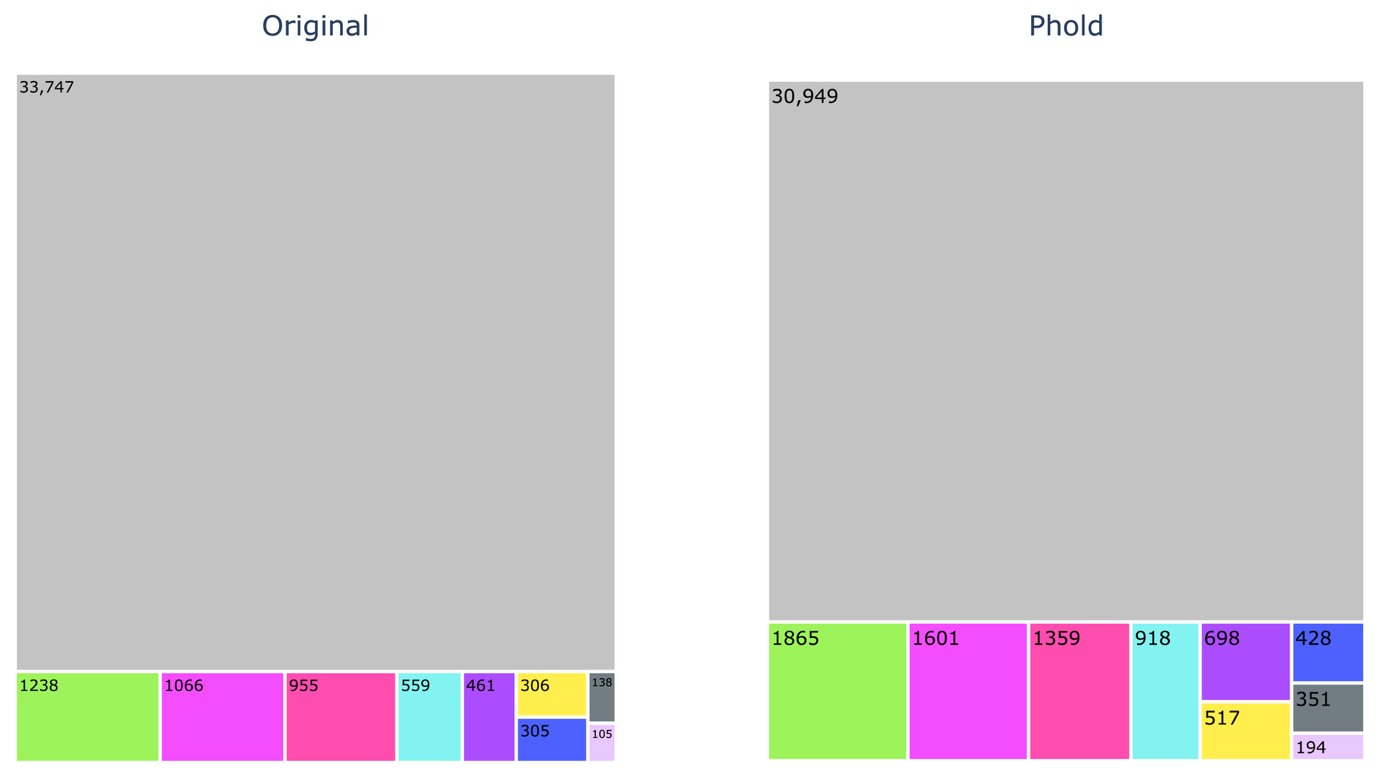

Supplementary Figure 6

Treemap plot showing the number of PHROG groups in each of the ten PHROG functional categories from the original PHROGs v4 database (left) compared to the Phold Search DB (right) with updated annotations. The colour scheme matches Figure 2b (i.e. in descending order of size on the left size, unknown function in grey, tail in green, nucleic acid metabolism in purple-pink, head and packaging in red-pink, other in light blue, moron AMG and host takeover in purple, transcription regulation in yellow, lysis in blue, connector in charcoal and integration and excision in pink).

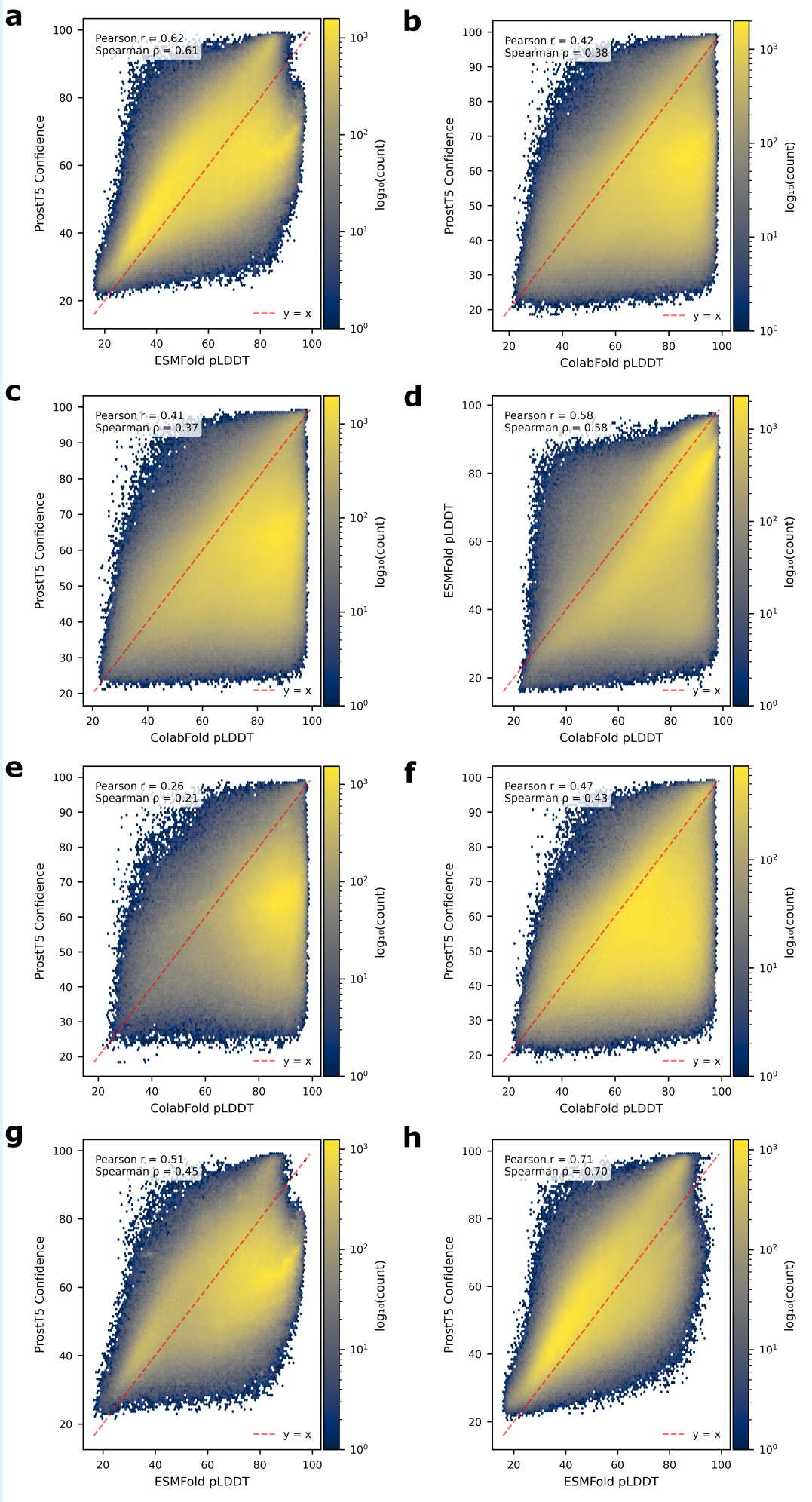

Supplementary Figure 7

ProstT5 confidence compared to various pLDDT. (a) Hexbin plot showing ProstT5 confidence (x-axis) and ESMFold pLDDT (y-axis) for all Phold DB 3.16M protein structures with ESMFold predictions. (b) Hexbin plot showing ProstT5 confidence (x-axis) and ColabFold pLDDT (y-axis) for all Phold DB 3.16M protein structures. (c) Hexbin plot showing ProstT5 confidence (x-axis) and ColabFold pLDDT (y-axis) for Phold DB 3.16M protein structures under 1000AA. (d) Hexbin plot showing ColabFold pLDDT (x-axis) and ESMFold pLDDT (y-axis) for all Phold DB 3.16M protein structures with ESMFold predictions. (e) Hexbin plot showing ProstT5 confidence (x-axis) and ColabFold pLDDT (y-axis) for all Phold Search DB 1.36M protein structures. (f) Hexbin plot showing ProstT5 confidence (x-axis) and ColabFold pLDDT (y-axis) for all 1.8M protein structures in Phold DB 3.16M without PHROG assignments. (g) Hexbin plot showing ProstT5 confidence (x-axis) and ESMFold pLDDT (y-axis) for all Phold Search DB 1.36M protein structures with ESMFold predictions. (h) Hexbin plot showing ProstT5 confidence (x-axis) and ESMFold pLDDT (y-axis) for all 1.8M protein structures in Phold DB 3.16M without PHROG assignments.

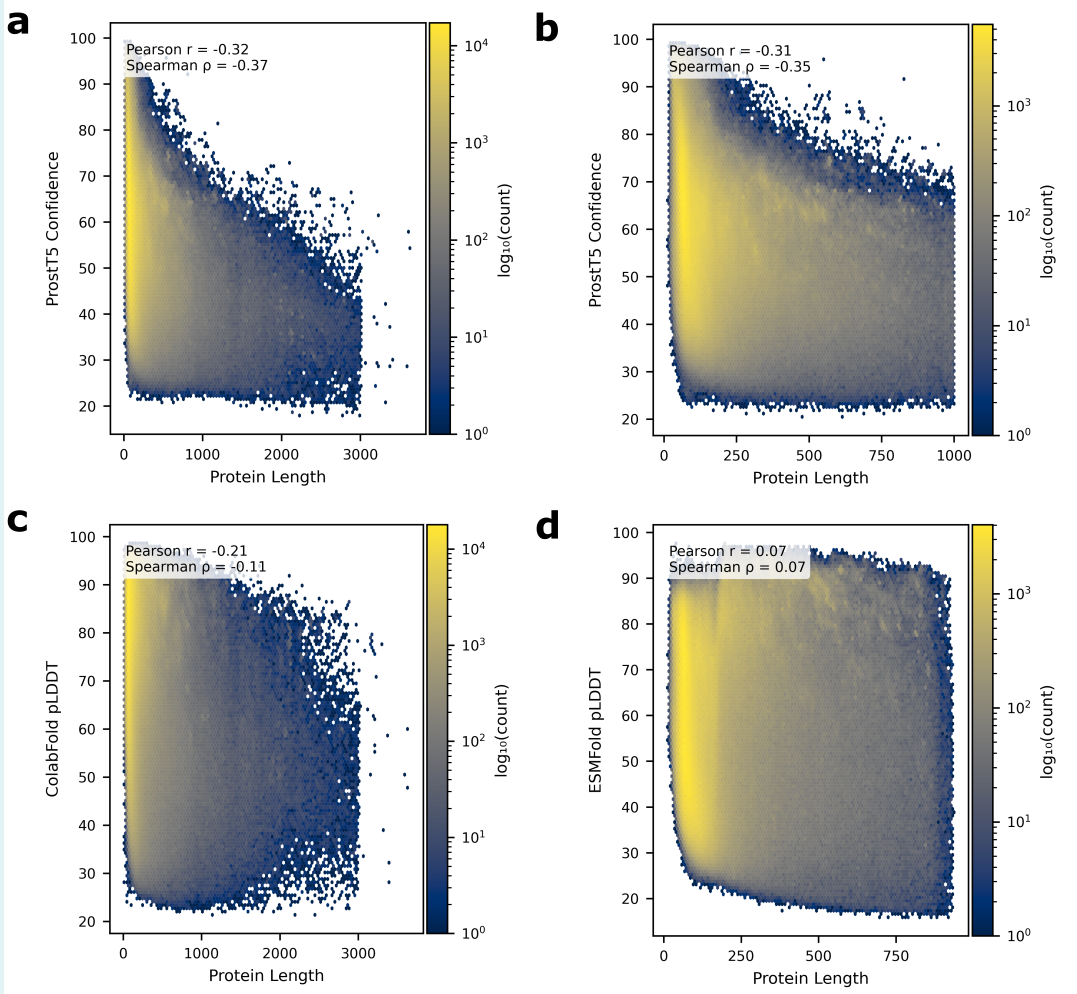
Supplementary Figure 8

ProstT5 confidence and pLDDT metrics compared to protein length. (a) Hexbin plot showing protein length (x-axis) and ProstT5 confidence (y-axis) for all protein structures in Phold DB 3.16M. (b) Hexbin plot showing protein length (x-axis) and ProstT5 confidence (y-axis) for all protein structures in Phold DB 3.16M under 1000AA. (c) Hexbin plot showing protein length (x-axis) and ColabFold pLDDT (y-axis) for all protein structures in Phold DB 3.16M. (d) Hexbin plot showing protein length (x-axis) and ESMFold pLDDT (y-axis) for all protein structures in Phold DB 3.16M with ESMFold predictions.

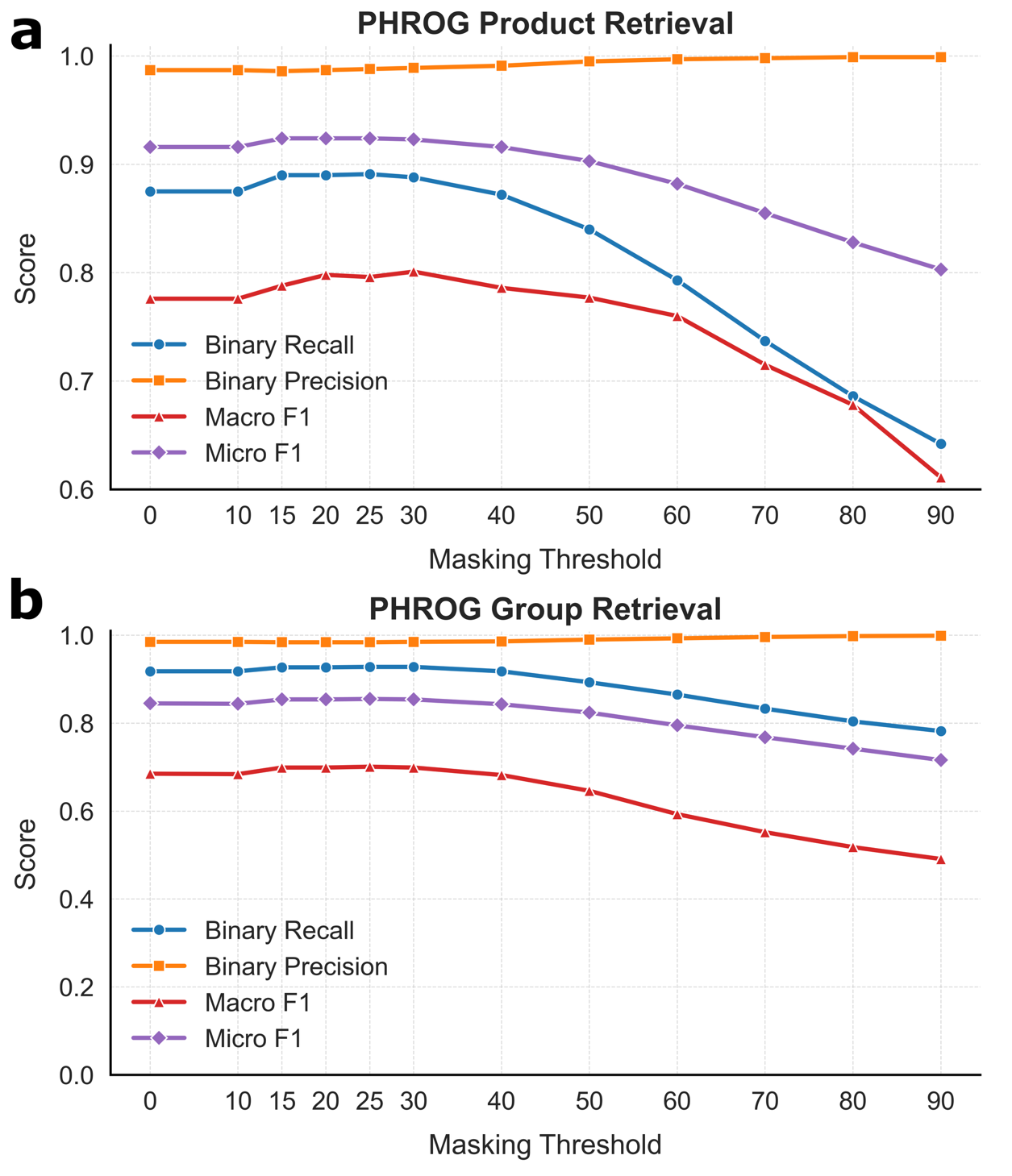

Supplementary Figure 9

PHROG annotation benchmarks when masking ProstT5 low confidence 3Di residues for different thresholds for the INPHARED 182 dataset for the (a) specific PHROG product level prediction task and (b) PHROG group prediction task. Note the y-axis is cropped and zoomed for (a) compared to (b).

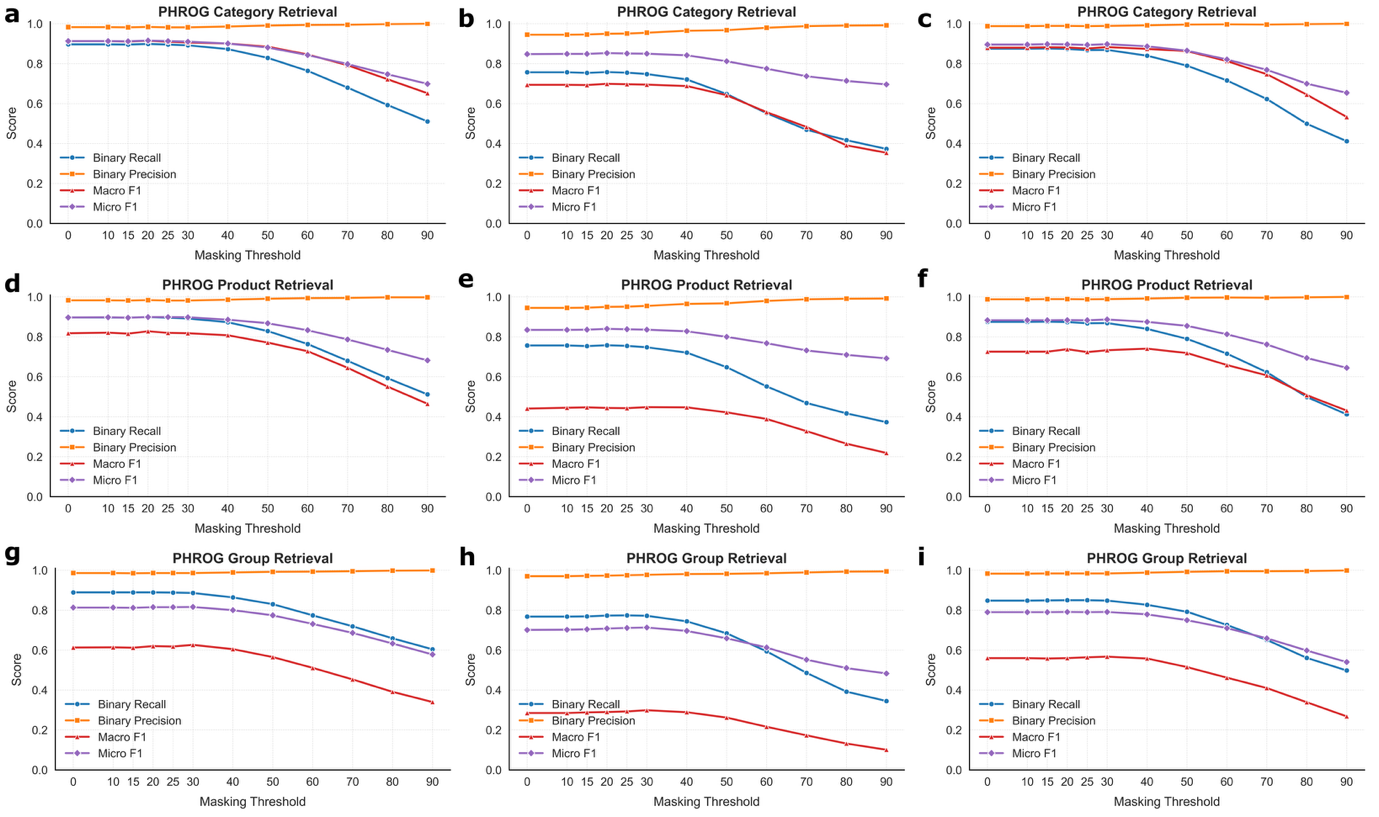

Supplementary Figure 10

PHROG annotation benchmarks when masking ProstT5 low confidence 3Di residues for different thresholds on the three metagenomic datasets. Metrics for the general PHROG category prediction task are presented for (a) Cook (b) Crass and (c) Tara datasets. Metrics for the specific PHROG product prediction task are presented for (d) Cook (e) Crass and (f) Tara datasets. Metrics for the PHROG group prediction task are presented for (g) Cook (h) Crass and (i) Tara datasets.

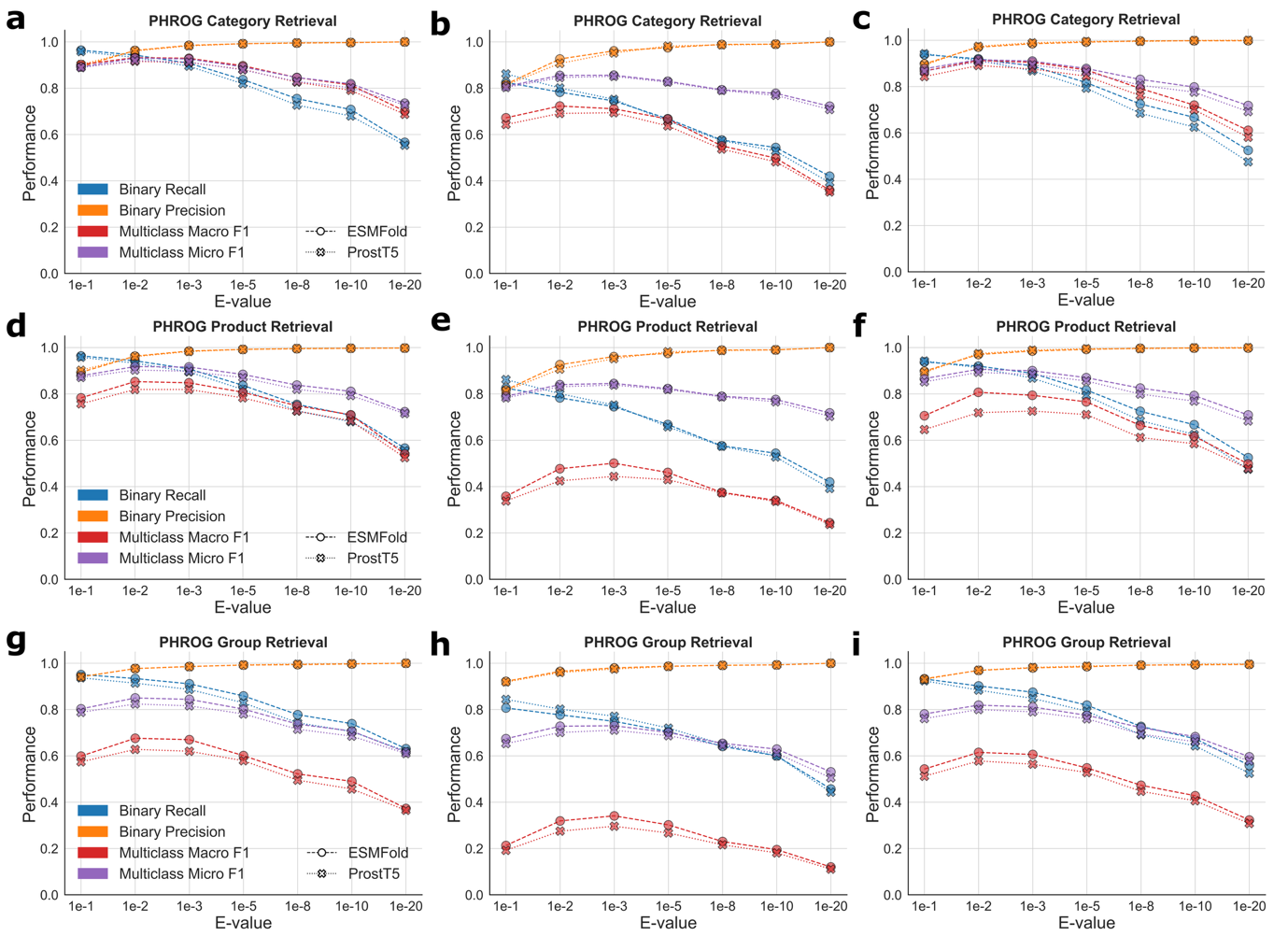
Figure 11

PHROG annotation benchmarks comparing Phold using ProstT5 and ESMFold for protein structure inference for different E-value thresholds. The PHROG category prediction task is shown for (a) Cook (b) Crass and (c) Tara, the specific PHROG product prediction task is shown for (d) Cook (e) Crass and (f) Tara and the PHROG group is shown for (g) Cook (h) Crass and (i) Tara datasets.

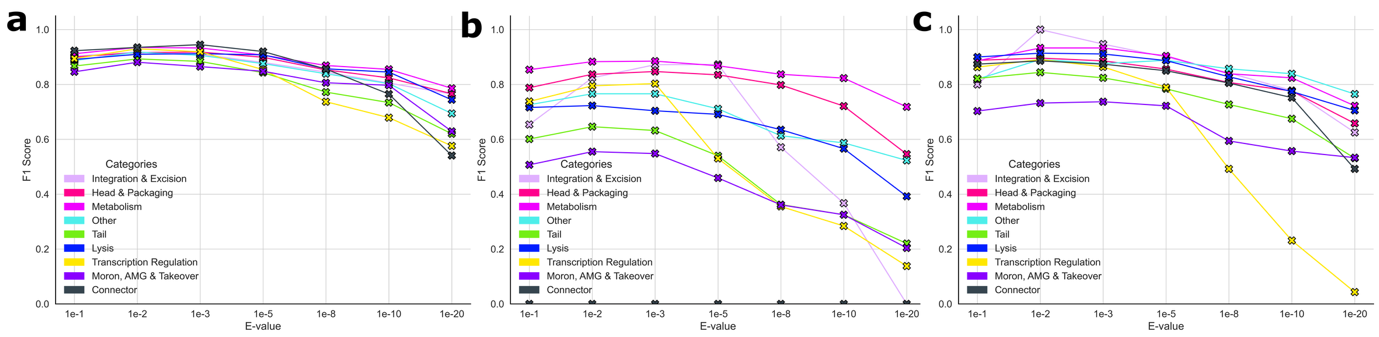

Supplementary Figure 12

Individual PHROG category annotation performance of Phold using ProstT5 at a variety of E-values on (a) Cook, (b) Crass and (c) Tara datasets compared to the pseudo ground truth of annotations with ColabFold generated structures at E-value 0.001. Note the F1-score for the connector category in (b) for Crass is 0 for every E-value.

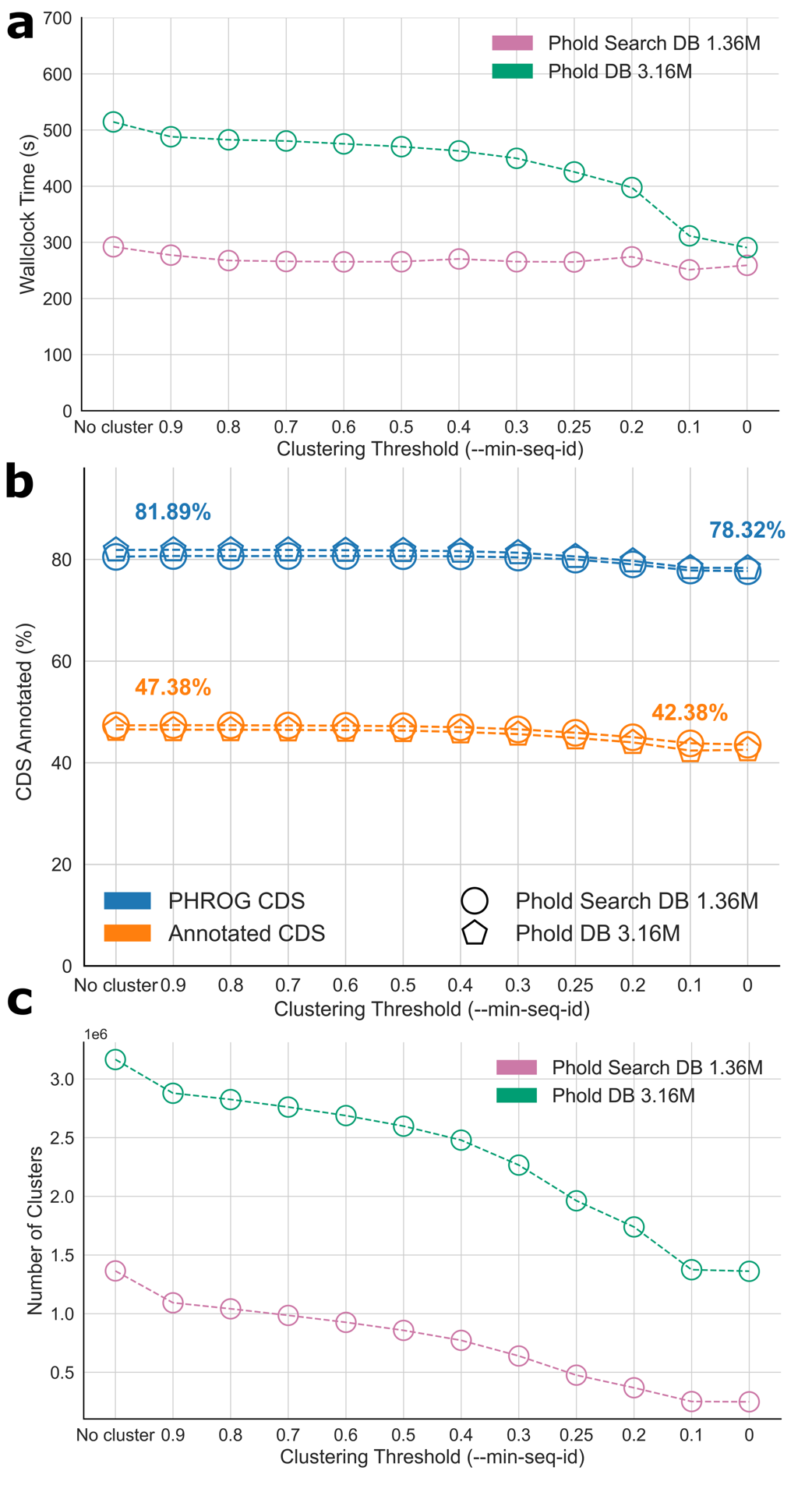

Supplementary Figure 13

Impact of database clustering on annotation performance. (a) Wallclock runtime in seconds (y-axis) of `phold compare` using pre-computed ProstT5 3Di predictions and Foldseek GPU on INPHARED 182 dataset using Phold with ProstT5 0/0/0000 0:00:00 amwith default parameters searching against Foldseek pre-clustered Phold Search DB 1.36M (purple) and Phold DB 3.16M (green) for different minimum sequence identity values (x-axis). (b) Functional (orange) and PHROG (blue, i.e. including unknown function PHROG) annotation rates with Phold using ProstT5 and Foldseek GPU across the 16,460 CDS in INPHARED 182 (y-axis) using the Phold Search DB 1.36M for different minimum sequence identity values (x-axis). (c) Number of Foldseek clusters (y-axis) for Phold Search DB 1.36M (purple) and Phold DB 3.16M (green) at different minimum sequence identity values (x-axis). The number of clusters are on the scale of 1e6.

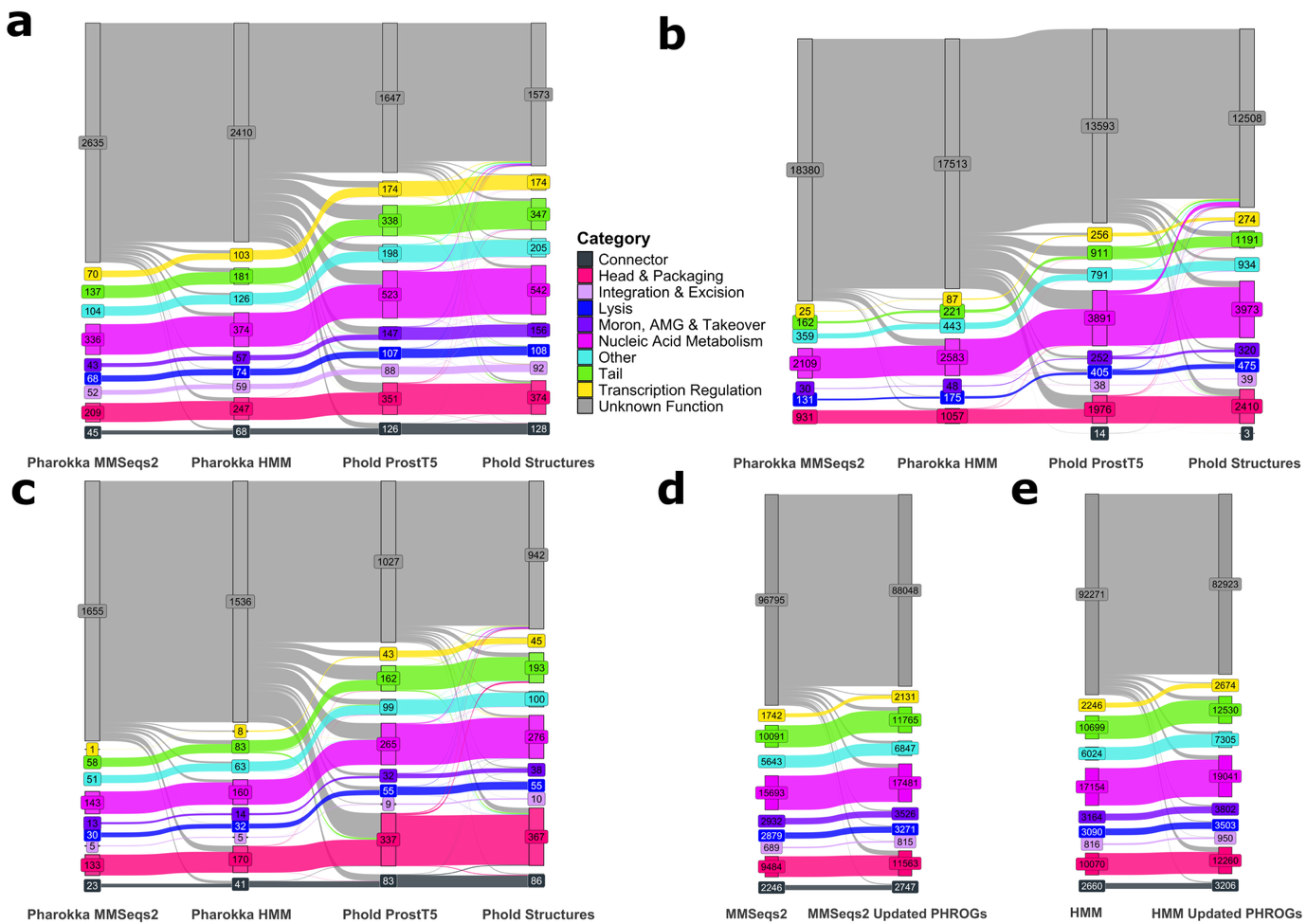

Supplementary Figure 14

Sankey plots showing: the flow of annotations between Pharokka with MMSeqs2, Pharokka with HMMs (PyHMMER), Phold with ProstT5 and Phold with structures for (a) Cook (b) Crass and (c) Tara datasets. Figure (d) shows the of annotations for the 1419 INPHARED phages between Pharokka with MMSeqs2 with the original PHROG v4 annotation labels (left) and the updated Phold annotation transfer labels with extra PHROG annotations manually curated as a part of this manuscript (right). Figure (e) shows the same as (d) but for Pharokka with PyHMMER.

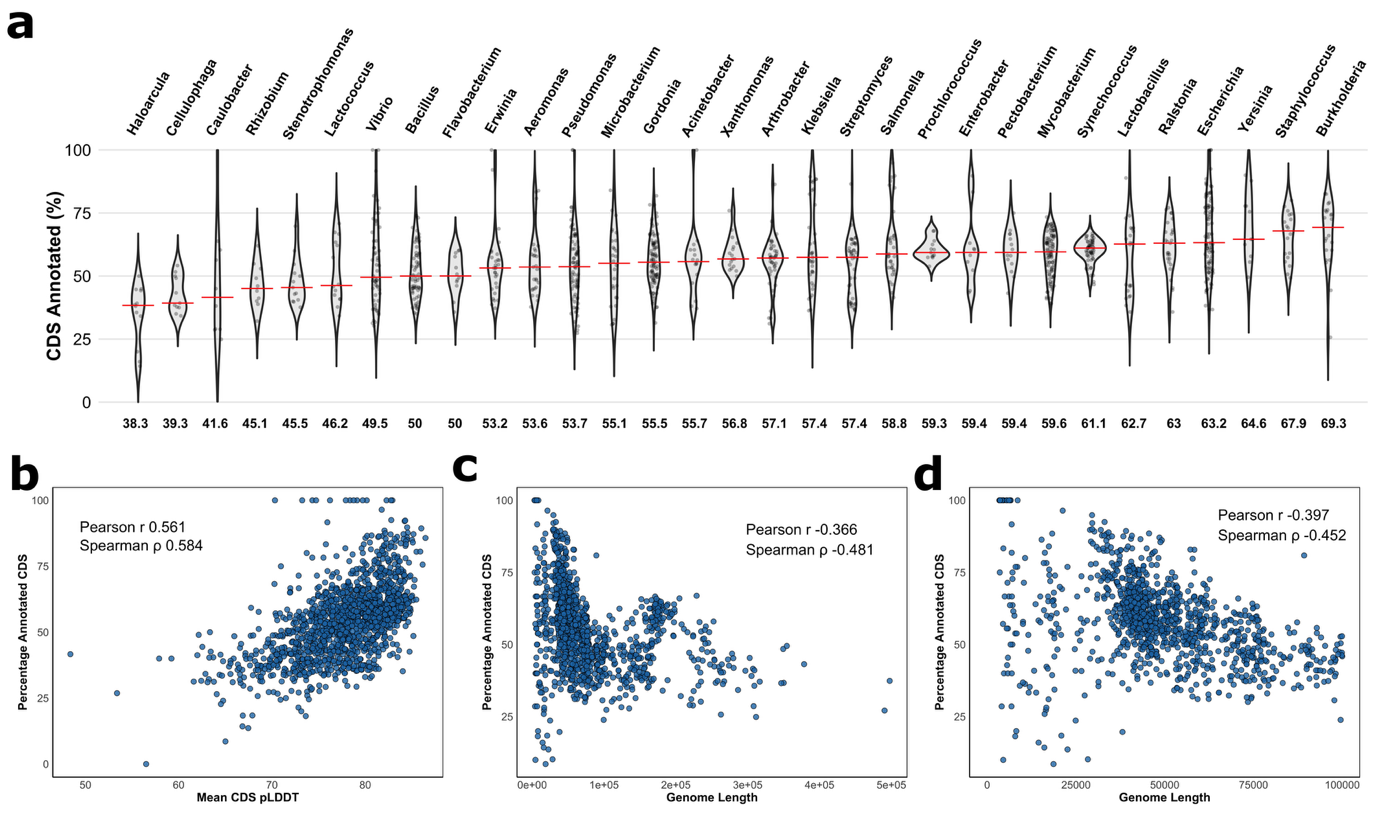

Supplementary Figure 15

Per-virus annotation rates in the INPHARED 1419 dataset. Figure (a) shows the percentage of annotated CDS using Phold with ColabFold structures for every virus in every host genus with at least 10 infecting viruses. The number below each violin plot is the median annotation rate for that host genus, also indicated on the plot with a red line. Figure (b) each point shows the mean CDS pLDDT across all CDS in the virus (x-axis) against the percentage of annotated CDS using Phold with ProstT5 in the virus (y-axis). Figures (c) and (d) show the viral genome length (x-axis) plotted against the percentage of CDS annotated using Phold with ProstT5 (y-axis). Figure (d) limits the x-axis to a maximum genome size of 100,000 bases.

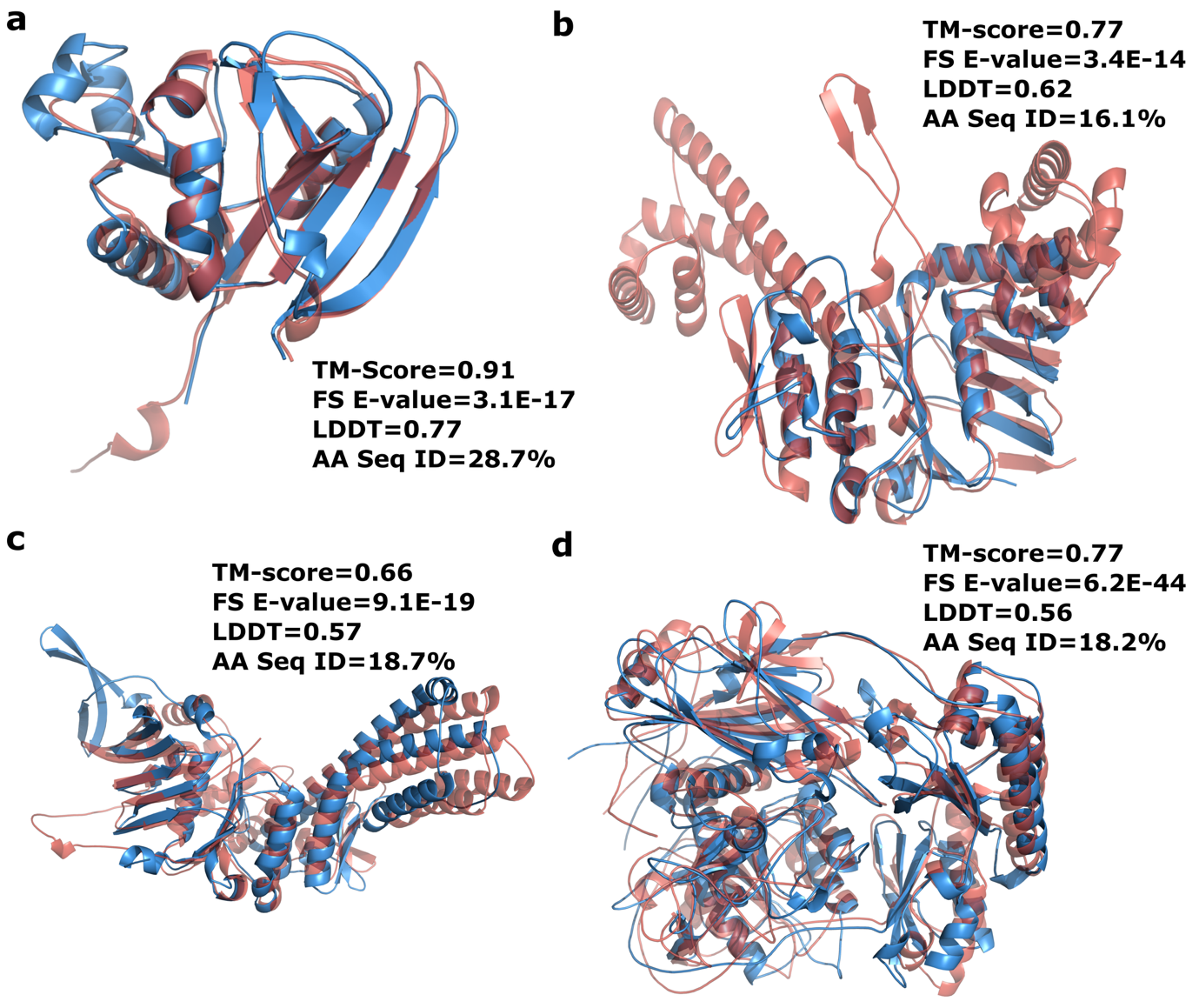

Supplementary Figure 16

ColabFold2 structure predictions comparing (a) the predicted dihydrofolate reductase from *Vibrio* phage VAP7 (GenBank accession: NC_048765) (blue) and CARD trimethoprim resistant dihydrofolate reductase gene dfrA26 (red); (b) predicted AriaA homolog to PARIS defense system subtype I on *Fusobacterium* phage vB_FnuS_FNU3 (GenBank accession: OQ808965); (c) predicted AriaA homolog to PARIS defense system subtype II on *Yersinia* phage vB_YenM_31.17 (GenBank accession: NC_070918) ; (d) and a letS protease homolog to the Gao_let defense system on *Enterocbacter* phage phiT5282H (GenBank accession: NC_049429).

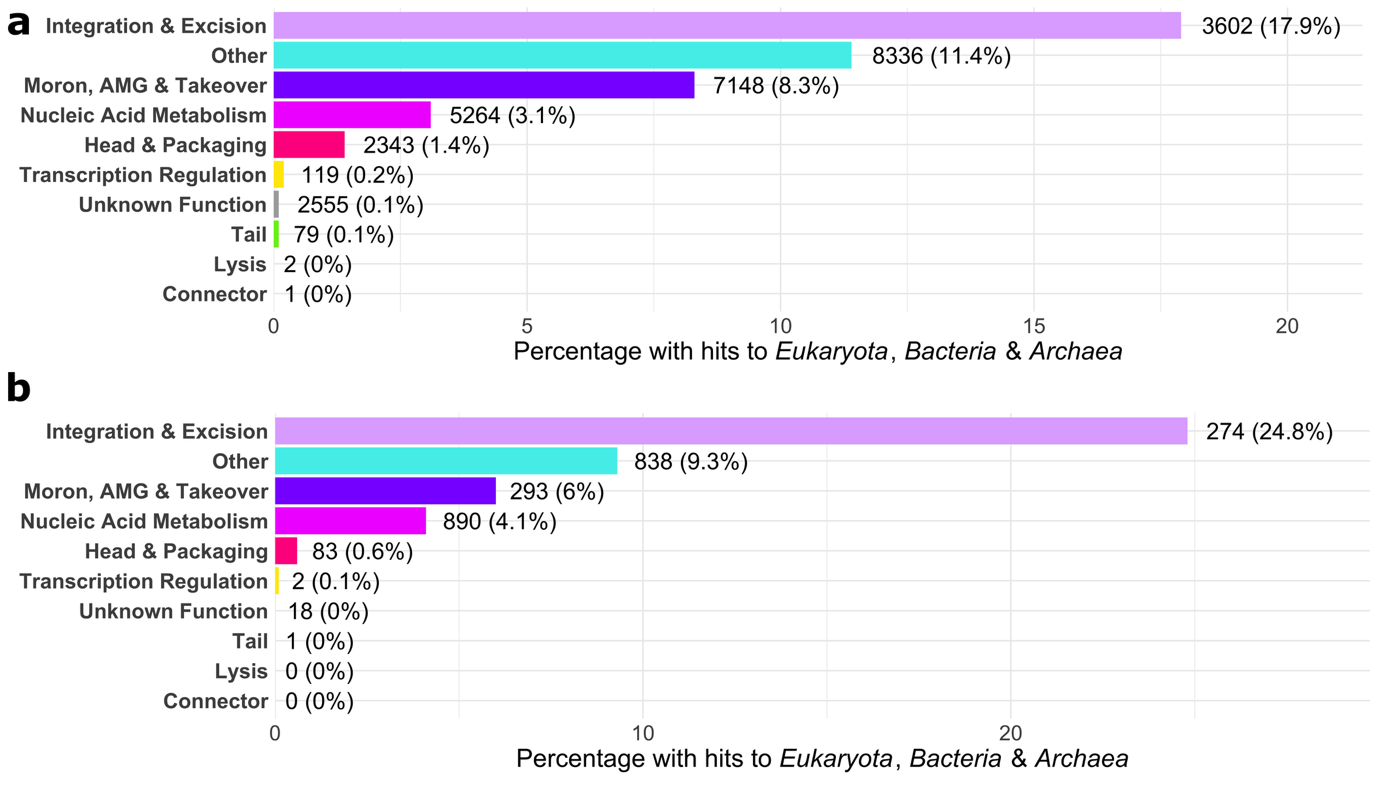

Supplementary Figure 17

Structural similarity between phage proteins across the tree of life: AlphaFold-Proteome. Percentage of proteins belonging to each PHROG category that have strong structural similarity (Foldseek E-value < 0.01 and alignment TM-score > 0.6) to all three domains of life (*Eukaryota*, *Bacteria,* and *Methanocaldococcus jannaschii*, the only representative of *Archaea*) from the 48 genomes comprising AlphaFold-Proteome database for (a) Phold DB 3.16M proteins and (b) INPHARED 1419 proteins. The raw number and percentage per category are indicated to the right of each bar.

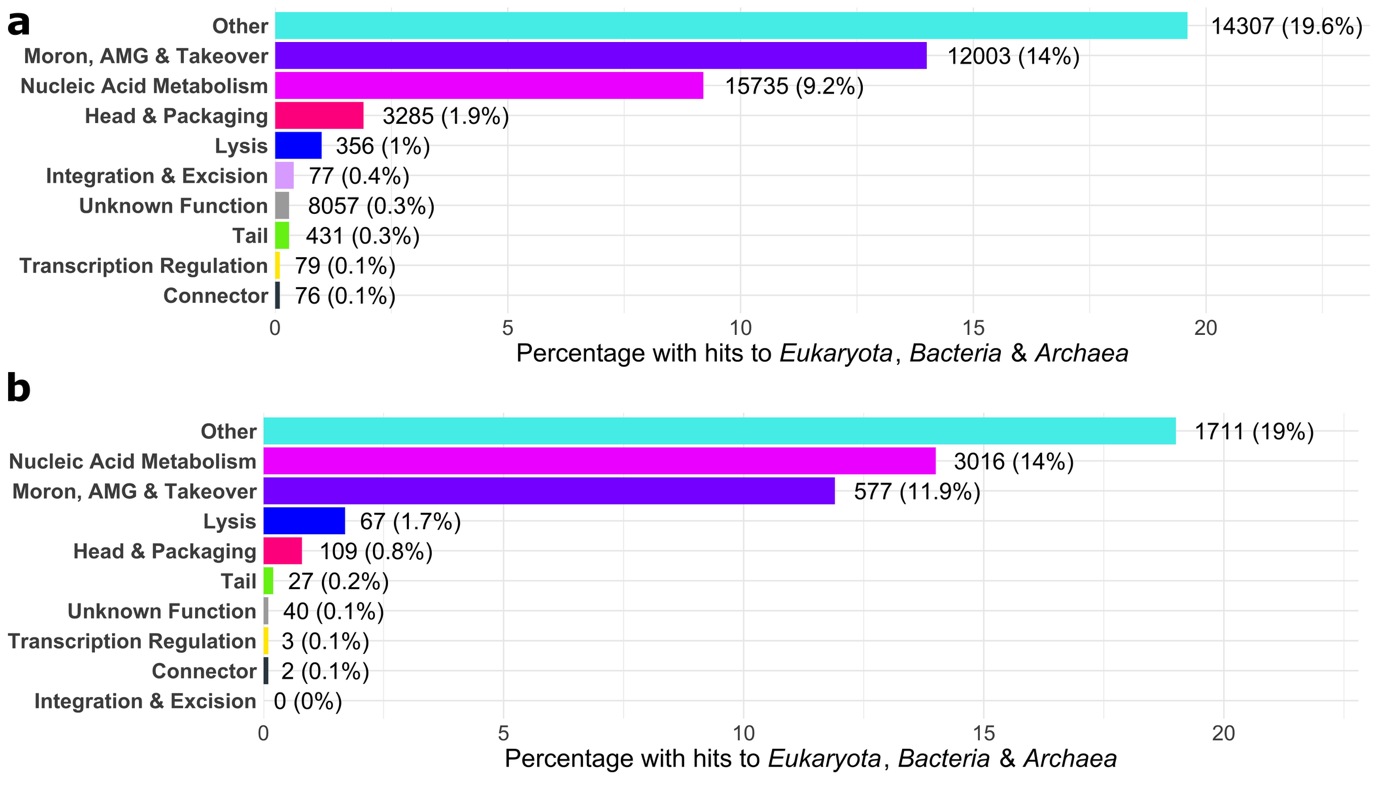

Supplementary Figure 18

Structural similarity between phage proteins to *Homo Sapiens* proteins. Percentage of proteins belonging to each PHROG category that have strong structural similarity (Foldseek E-value < 0.01 and alignment TM-score > 0.6) to *Homo sapiens* for (a) Phold DB 3.16M proteins and (b) INPHARED 1419 proteins. The raw number and percentage per category are indicated to the right of each bar.
