## Supplementary Notes for "Protein Structure Informed Bacteriophage Genome Annotation with Phold"

#### Supplementary Note 1: ColabFold and ESMFold Protein Structure Prediction Comparisons

For Phold DB 3.16M, we were able to predict protein structures with both ColabFold^1^ (implementing Alphafold2^2^) and ESMFold^3^ for 3,041,605, retaining the prediction with higher pLDDT for Phold. ColabFold-only predictions were generated for the remaining 124,597 due to ESMFold GPU VRAM constraints. The ColabFold prediction had the higher pLDDT for 2,752,972/3,166,202 (86.5%) proteins while the ESMFold prediction was selected for 413,230 (13.5%). ColabFold structures overall had higher pLDDT (mean: 74.80) than ESMFold structures (mean: 61.18). This pLDDT difference was larger for the 1,802,498 enVhog and efam proteins not assigned to PHROGs (ColabFold mean: 70.16 vs ESMFold: 54.78) compared to all 1,319,865 proteins assigned to PHROGs (ColabFold mean: 80.78 vs ESMFold: 69.19). There was a positive relationship between both MSA depth (measured by the number of homologs) and quality (NEFF^4^) with pLDDT for ColabFold (Spearman’s ρ: 0.33 and 0.41 respectively) and ESMFold (Spearman’s ρ: 0.34 and 0.16 respectively) (Figure 2C, Supp Figure 2A). For ColabFold, there was little difference between PHROGs and the less well-characterised enVhog, efam and singleton proteins (PHROGS only: homologs vs pLDDT Spearman’s ρ: 0.33 Supp Figure 2B; efam, enVhog and singletons: Spearman’s ρ: 0.30 Supp Figure 2C). As shown previously, the performance of pLM-based predictors correlates well with the number of sequence homologs in today’s databases, presumably because seeing more examples of the same/similar fold helps learning better representations^3^ and because pLMs implicitly learn co-evolutionary information that informs structure during pre-training^5^. Accordingly, ESMFold pLDDT improved only when the number of homologs was at least 100, and substantially so only above 500 (Figure 2C). For proteins with fewer than 100 homologs, there was no correlation between ESMFold pLDDT and number of homologs (Spearman’s ρ: -0.01, median pLDDT: 54.6), but for those with at least 100 homologs, there was a positive correlation and more accurately predicted structures (Spearman’s ρ: 0.30, median pLDDT: 67.6).

In terms of PHROG functional categories, there was a greater difference between pLDDT for ColabFold and ESMFold predictions for unknown function PHROG proteins (mean pLDDT difference 16.03) compared to other categories (pLDDT differences from 5.86 for integration and excision to 12.18 for moron, AMG and takeover) (Supp Table 1). 815/856 unique PHROG product descriptions had mean pLDDT of at least 70 (Supplementary Table 2). Excluding five product descriptions for which no ESMFold predictions could be generated, 806/851 had higher mean pLDDTs for ColabFold-generated structures compared to ESMFold-generated structures. 6/10 PHROG protein product descriptions with the largest difference where ESMFold had higher mean pLDDT than ColabFold were in the tail category, including ‘tail length tape measure protein’ (14,846 proteins) and ‘pilot protein for DNA ejection’ (465 proteins).

### Supplementary Note 2: LoRA Finetuning

Amidst our analyses of ProstT5 confidence and accuracy, we also finetuned the full ProstT5 encoder-decoder model using the low-order rank adaption (LoRA) approach and trained CNNs on top of the finetuned embeddings, aiming to improve ProstT5’s performance on phage proteins. While our LoRA finetuned ProstT5 encoder-decoder model improved 3Di reconstruction accuracy compared to the standard ProstT5 encoder-decoder on the validation set of phage proteins from 64.5% to 68.3%, the improvement was small for the finetuned encoder plus CNN models (improving 3Di reconstruction from 63.6% to 64.5%) (Supplementary Figure 5). The differences in downstream annotation performance for the finetuned encoder plus CNN were negligible compared to standard ProstT5 with CNN (Supplementary Table 9). Given the unfeasible computational demands of the encoder-decoder for 3Di inference^6^ (i.e. the full encoder-decoder model is larger (requiring larger GPUs for inference) and far slower; generating 3Di predictions using the full encoder-decoder for 1344 Phold DB validation proteins took 53 minutes on an NVIDIA A100 40GB GPU, compared to 36 seconds for the encoder plus CNN) and the lack of improvement in the finetuned encoder with CNN mode we chose to keep the original ProstT5 encoder and CNN for Phold.

#### Supplementary Note 3: ProstT5 Confidence Analysis

ProstT5 confidence was more strongly correlated with ESMFold pLDDT (Spearman’s ρ 0.61 (Supp Figure 6A), than ColabFold pLDDT (Spearman’s ρ 0.38 Supplementary Figures 7B-C), even when considering only proteins below 1000AA, possibly reflecting the shared pLM underpinnings of ProstT5-CNN and ESMFold. ESMFold was in fact more strongly correlated with ProstT5 confidence than ColabFold pLDDT (Pearson 0.58, Spearman’s ρ 0.58) (Supplementary Figure 7D). Proteins not assigned to PHROGs were more strongly correlated with both ESMFold and ColabFold pLDDTs than those in the Phold Search DB (Supplementary Figures 7E-H). ProstT5 confidence was negatively correlated with protein length, particularly showing higher values for shorter proteins below 100AA, likely explaining the higher correlation with proteins not assigned PHROGs, as these are shorter on average than those that are (Supplementary Figures 8A-B). In contrast, ColabFold pLDDT showed a slight negative correlation whereas ESMFold pLDDT was slightly positively correlated with length (Supplementary Figure 8C-D).

### Supplementary Note 4: Phold Database Clustering Ablations

We also compared Phold’s annotation performance using the Phold Search DB 1.36M, consisting of only proteins with an assigned PHROG along with the other specialised databases, to using Phold DB 3.16M that also includes enVhog and efam proteins with no PHROG assignment. We concurrently ran ablations to determine whether pre-clustering the databases at a variety of minimum sequence identity values and running Foldseek searches against the cluster representatives then expanding this to all members would provide a better trade-off in terms of annotation performance against runtime.

Overall, applying the default Foldseek sensitivity of `--max-seqs 1000, pre-clustering at low levels of sequence similarity (i.e. increasing the tightness of clustering) reduced annotation performance (Supplementary Figure 13). Runtimes were similar across all clustering values, and higher when using the Phold DB 3.16M than the Phold Search DB 1.36M. Using the larger Phold DB 3.16M enabled more PHROG group annotations than the Phold Search DB 1.36M, though there were negligible differences in functional annotation rates. This indicates that using the larger database finds more hits to unknown function proteins, but not functionally annotated proteins, and may be better suited to identifying novel protein that are likely phage-like, rather than more detailed annotation tasks. We chose to implement the Phold Search DB 1.36M as the default search database in Phold, given reduced runtime and similar functional annotation rates.
